# BRAIN CAST: An MRIQC-guided pipeline for age- and sex-specific pediatric brain MRI template construction, validated by downstream structural fidelity

**DOI:** 10.64898/2026.08.02.742256

**Authors:** Yang Hu, Jose Contreras-Vidal

## Abstract

Pediatric neuroimaging needs age- and sex-appropriate references, yet existing atlases span broad age ranges that blur development or lack sex specificity. We present BRAIN CAST: 28 year-by-year, sex-specific brain MRI templates covering ages 5–18, built from 1,272 quality-screened children in the Healthy Brain Network by an MRIQC-guided pipeline combining reduced-strength denoising, cerebrospinal-fluid–anchored intensity normalization, deep-learning skull stripping and iterative groupwise diffeomorphic registration. We evaluate templates not by image sharpness, which is not comparable across intensity conventions, but by the structural bias they induce downstream. Held-out children align to their matched template with sub-voxel gray–white interface error (1.1 mm); on a direction-symmetric surface-distance metric BRAIN CAST matches the best single-template reference and outperforms an age-specific pediatric atlas in 189 of 189 subjects. Female cortex is fit measurably better by female than by male templates, an effect no sex-neutral reference can provide. Templates, tissue-probability maps and the containerized pipeline are released.

## 1 Introduction

The human brain undergoes a period of profound and dynamic development throughout childhood and adolescence, characterized by rapid and nonlinear changes in structure, function, and organization [8–10]. Understanding these developmental trajectories, which include processes like cortical thinning, white-matter myelination, and ventricular expansion, is crucial for mapping typical brain maturation and identifying atypical patterns associated with neurodevelopmental and psychiatric disorders. Magnetic Resonance Imaging (MRI) has become an indispensable tool in this endeavor, providing non-invasive, in-vivo measurements of brain anatomy and function. However, the accuracy and reliability of pediatric neuroimaging studies depend critically on the use of appropriate age- and population-specific analytical tools, particularly brain templates.

### 1.1 The Need for Pediatric-Specific Brain Templates

Pediatric brains are not simply scaled-down versions of adult brains. Several key factors necessitate the use of dedicated pediatric brain templates. First, the brain undergoes **rapid and nonlinear neurodevelopment** from childhood to early adulthood [8]. This includes significant changes in cortical thickness, myelination, and the size and shape of subcortical structures [8, 9, 11]. Second, the **tissue properties** of the developing brain differ from those of the mature brain. For instance, ongoing myelination results in lower gray matter–white matter (GM–WM) contrast in school-age children, and the cortical ribbon is thinner with evolving folding patterns [9]. Third, **sex differences** in brain structure emerge and become more pronounced during adolescence, with subtle but consistent variations in the shape and volume of the cortex and ventricles [12–14]. Using adult templates or even age-inappropriate pediatric templates can introduce significant biases, large registration errors, and intensity mismatches, which can obscure real developmental effects or generate spurious group differences [6, 15, 16].

### 1.2 The Importance of High-Fidelity Pediatric Anatomy

High-fidelity, age- and sex-matched pediatric brain templates are essential for a wide range of neuroimaging applications. With an accurate, population-specific reference, image registration becomes more precise, improving the sensitivity of voxelwise statistical analyses and morphometric measurements. For example, sharper templates with well-defined structural boundaries yield better inter-subject alignment, which in turn enhances the detection of subtle growth or atrophy patterns. This is especially important for surface-based analyses, as the pediatric cortex is approximately 30% thinner than that of adults and is prone to misclassification if anatomical edges are blurred. In **structural MRI analysis**, accurate spatial normalization to a representative template is a prerequisite for precise morphometric measurements [7]. For **functional MRI (fMRI)**, proper alignment of cortical folding patterns is critical for group-level activation mapping. In the context of **electroencephalography (EEG) and magnetoencephalography (MEG)**, the accuracy of source localization depends on the geometric accuracy of the forward head model. The use of adult or mismatched head models has been shown to result in large localization errors, on the order of 10–20 mm, while age- and sex-appropriate templates significantly improve dipole localization accuracy [17, 18]. The utility of such references is further amplified when they are distributed as standardized, redistributable reference spaces with consistent naming conventions [19] and shared through open, FAIR-native archives that maximize reuse of derived neuroimaging resources [20].

### 1.3 Gaps in Existing Methodologies

Despite the clear need for high-quality pediatric brain templates, existing resources have significant limitations in age coverage, cohort size, and demographic specificity. Many publicly available atlases average across **broad age ranges**, diluting age-specific features. For instance, the NIH pediatric atlas spans from 4.5 to 18.5 years in a single continuous template, an approach that overlooks rapid developmental changes [7]. Conversely, recent dense, age-resolved (4D) pediatric atlases concentrate on infancy and the first years of life, such as the Baby Connectome Project volumetric atlas covering 0–24 months [21], leaving the school-age and adolescent 5–18 range comparatively underserved. Some templates are derived from relatively **small sample sizes** (tens of subjects), raising concerns about noise and population bias. Even comprehensive age-specific resources such as the Neurodevelopmental MRI Database, which spans childhood into adolescence, are assembled by pooling scans from multiple prior studies rather than from a single, uniformly screened cohort [6]. Furthermore, most prior templates do not explicitly address **sex differences** in brain development. While some population-specific templates exist, such as the Chinese pediatric atlas (CHN-PD) from Beijing Normal University, they cover only a narrow school-age range (6–12 years) and are limited to their source population [22]. In summary, the field has lacked a comprehensive set of pediatric templates with fine-grained age resolution and consideration of sex, built on a sufficiently large and heterogeneous dataset.

### 1.4 The Present Study

To address these critical gaps, we present a new set of high-resolution, age- and sex-specific pediatric brain MRI templates spanning ages 5 through 18 years. We leveraged the large-scale Healthy Brain Network (HBN) initiative—a community pediatric neuroimaging biobank centered in the New York City area—as our primary data source [1]. We selected HBN because, among openly available pediatric datasets, it uniquely pairs a large single-initiative sample spanning the full 5–18 school-age range on harmonized 3 T protocols with sufficient per-year, per-sex subject counts to build fine-grained age-and sex-specific templates—coverage that neonatal/infant resources and smaller or narrower-age atlases do not provide. Over 2,400 T1-weighted MRI scans from a community pediatric cohort were aggregated and screened for gross structural abnormality and image quality, with 1,272 contributing to the final templates—an unprecedented sample size for pediatric template construction. By stratifying the data into narrow one-year age bands and further by sex within each year, our templates closely track the subtle anatomical changes that occur from early childhood into late adolescence. This design yields a library of developmental brain templates, each reflecting a specific age and sex subgroup with up to roughly 170 subjects per year-group, ensuring both high anatomical detail and robust population representativeness.

### 1.5 Methodological Innovations

We also introduce several methodological innovations in the template construction pipeline to address pediatric-specific imaging challenges. First, we apply spatially adaptive non-local means (SANLM) denoising to suppress noise while preserving the fine tissue contrasts characteristic of developing brains [23]. Second, for skull stripping, we employ a deep learning–based brain extraction tool, DeepBET, which uses a 3D U-Net to reliably isolate brain tissue [4]. Third, we perform intensity normalization anchored on cerebrospinal fluid (CSF) signal, using the ventricles as a stable reference to align intensity scales across subjects. Finally, we construct the templates using a diffeomorphic (smooth, invertible, and topology-preserving) registration algorithm with a symmetric normalization (SyN) transformation integrated in the Advanced Normalization Tools (ANTs) framework [5, 24]. We disabled the usual iterative intensity rescaling in template building to preserve the original tissue contrast and adjusted regularization parameters to account for the thinner cortex and higher anatomical variability in youth.

### 1.6 Template Validation

We quantitatively evaluated the new templates against existing standards to validate their quality. Objective image quality metrics were used, including an edge sharpness index to assess the clarity of anatomical boundaries [25] and gray–white matter contrast-to-noise ratios to gauge tissue separability. We also examined the deformation fields required to warp individual brains to each template, reasoning that a better template would require smaller and smoother deformations on average. Across these evaluations, the templates constructed in this work showed anatomical fidelity comparable to a contemporary single-template reference (NKI) on registration-based metrics, decisively exceeded a broad-age pediatric reference (Fonov/NIHPD) in downstream structural fidelity, and provided fine-grained age and sex specificity together with higher scale-invariant structural detail—contributions that a single sex-neutral, broad-age reference cannot offer.

## 2 Methods

### 2.1 Constraint-aware Pediatric Preprocessing

Standard neuroimaging pipelines optimized for adult brains often falter when applied to pediatric MRI, due to stark developmental differences in brain size, contrast, and morphology [2, 26]. We addressed these issues by treating preprocessing as a constrained optimization problem that strictly preserves key anatomical features (ventricular volume, cortical topology). In particular, brain extraction and tissue segmentation steps were tuned to avoid overmasking (excess removal of developing cortex or enlarged ventricles) and undermasking (retaining non-brain matter that could corrupt intensity distributions). Age-specific brain atlases [26] and pediatric tissue probability maps were incorporated to guide these steps, ensuring that priors reflect age-appropriate ventricular size and gray/white matter contrast. Where adult templates would misalign or misclassify regions in children, our pipeline uses iterative atlas refinement to minimize such errors [2]. This constraint-aware approach maintained anatomical validity by imposing morphological priors (e.g. smoothness constraints that do not distort ventricular geometry) during optimization. As a result, each preprocessing module (skull stripping, bias correction, etc.) was parameterized to prioritize the retention of true anatomy over blind artifact removal. Known failure modes of adult pipelines (e.g. skull strippers cutting into pediatric cortex or merging CSF with gray matter) were mitigated through these tailored constraints, yielding cleaner brain masks and more reliable tissue classification for young children. In summary, the pediatric preprocessing was formulated as an optimization under anatomical constraints, explicitly balancing aggressiveness in artifact cleanup with fidelity to age-specific brain morphology.

#### Preprocessing stage (Steps 1–10)

Preprocessing consisted of ten ordered steps designed to preserve pediatric anatomical fidelity while improving inter-subject comparability for downstream registration and template construction (Fig. 1).

**Fig. 1:**
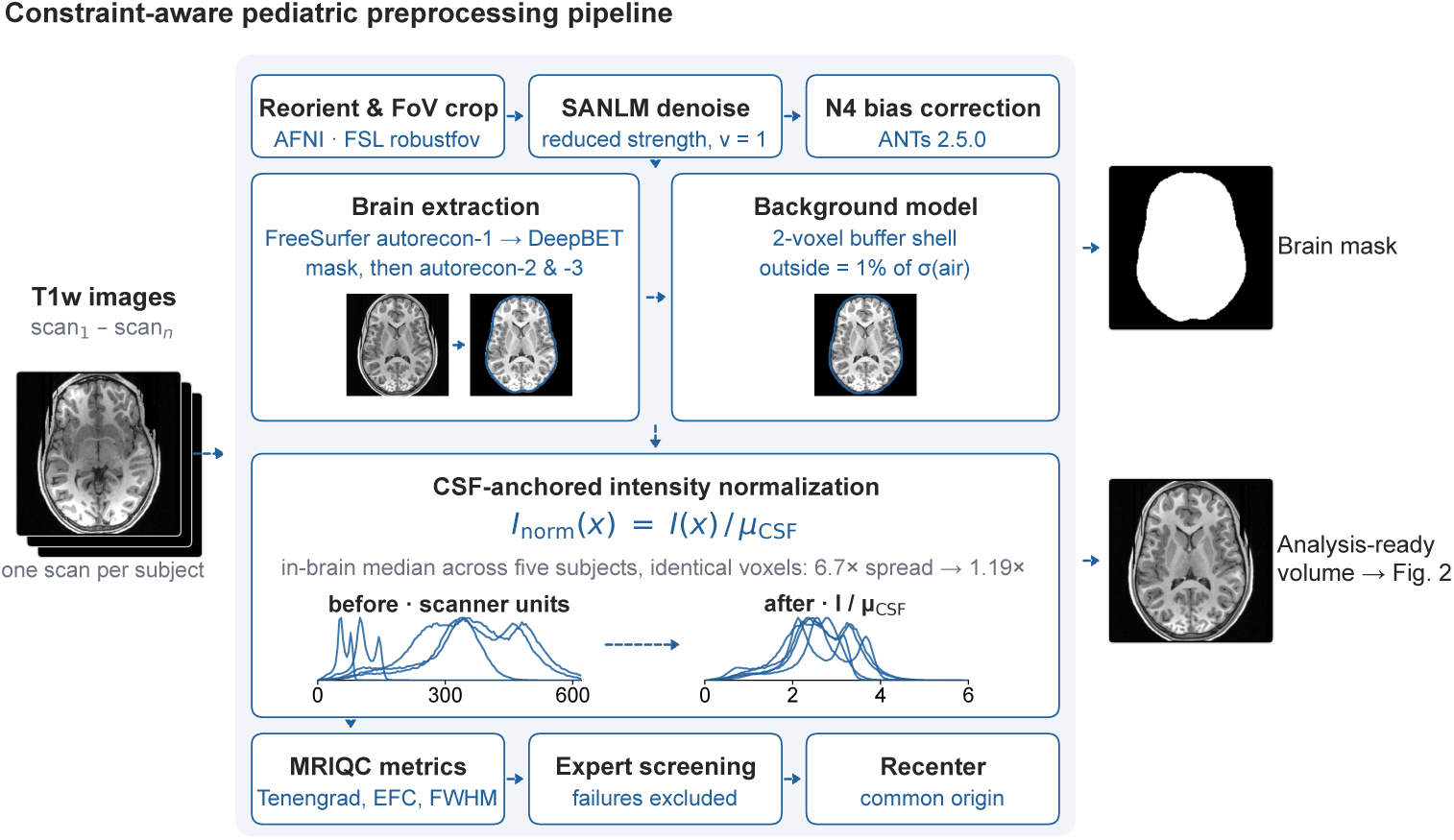
Constraint-aware pediatric preprocessing pipeline. Raw Healthy Brain Network T1-weighted scans are conditioned, brain-extracted and placed on a common intensity scale before template construction; tools and versions are given per step, and parameter choices are described in the Methods. Every brain panel is a real intermediate volume from one subject’s run of the released pipeline, shown at the same axial slice throughout. The pediatric-specific decisions are the reduced-strength SANLM filter (*v* = 1), the hybrid FreeSurfer-plus-DeepBET brain mask (outlined), the explicit background model — a two-voxel buffer shell around the brain (outlined), with every voxel beyond it set to 1% of the measured air-background standard deviation — and the CSF-anchored intensity scale. Because that scaling divides a volume by a single number, it cannot alter a subject’s own contrast; its purpose is to make different subjects commensurable, and the panel measures exactly that. In-brain intensity distributions of five subjects, computed over identical voxels before and after, show the between-subject spread of the median intensity falling 6.7-fold to 1.19-fold. The resulting scale is carried through template construction unchanged (Fig. 2).

#### Step 1: Reorientation and field-of-view cropping

All scans were reoriented to a common RPI orientation (right-to-left, posterior-to-anterior, inferior-to-superior voxel-axis ordering) using AFNI’s 3dresample, providing a consistent axis convention for downstream ANTs and FSL tools, and then subjected to robust field-of-view cropping (FSL robustfov) to remove non-brain regions, including neck and lower head structures, while preserving inferior and posterior brain anatomy. This step standardizes image orientation and spatial extent across subjects without truncating cerebellar or occipital regions, thereby improving consistency for subsequent processing and registration.

#### Step 2: Conservative denoising

Spatially adaptive non-local means (SANLM) denoising was applied using a reduced filtering strength. SANLM accounts for spatially varying noise distributions commonly observed in MRI and has been shown to outperform global smoothing approaches in preserving anatomical detail [23]. While aggressive denoising is often used in adult pipelines to maximize signal-to-noise ratio, such settings can blur thin cortical ribbons and low-contrast tissue boundaries in pediatric brains. We therefore used a conservative parameterization to attenuate thermal noise while preserving fine-scale gray–white matter contrast critical for accurate segmentation and registration.

#### Step 3: Bias-field correction

Intensity inhomogeneity was corrected using the N4 bias-field correction algorithm [27]. Unlike many adult pipelines that apply bias correction after skull stripping, N4 was applied prior to brain extraction. In pediatric MRI, smooth intensity gradients induced by coil sensitivity or head-size variability can mimic tissue boundaries and compromise skull stripping at thin cortical margins. Performing N4 correction on the full image reduces such confounds, ensuring that subsequent brain extraction is driven by anatomical rather than scanner-related intensity variation.

#### Step 4: Surface-based anatomical reconstruction

An initial anatomical reconstruction was performed using the surface-based processing stream of FreeSurfer [28]. FreeSurfer autorecon estimates white matter and pial surfaces through topology-constrained deformable models, providing anatomically informed priors on brain boundaries. In pediatric data, surface-based approaches are less prone to gross over-stripping than purely intensity-based methods, particularly in regions with thin cortex or enlarged ventricles.

#### Step 5: Background noise modeling

To preserve realistic image statistics, background voxels were explicitly modeled rather than zeroed. Following initial masking, a narrow buffer shell surrounding the brain was retained at its original intensities, and all voxels beyond that shell were set to a small constant scaled to the measured air-region standard deviation rather than to zero. This strategy creates a smooth intensity transition at the brain–air interface, reduces resampling artifacts, and maintains a non-zero floor. A non-zero background is required in practice by FreeSurfer’s recon-all, which fails on a fully zeroed brainmask; it also keeps automated quality metrics such as the foreground–background energy ratio (FBER) and entropy focus criterion (EFC) well defined, since these become ill-behaved when background intensities are artificially set to zero [29].

#### Step 6: Deep learning–based brain masking

Brain extraction was refined using DeepBET, a fast three-dimensional convolutional neural network for skull stripping [4]. DeepBET employs a two-stage architecture based on LinkNet [30] and U-Net [31], in which an initial coarse mask is used to crop the image before predicting a refined final brain mask. Although trained primarily on adult T1-weighted MRI, DeepBET has been shown to generalize well across anatomical variability, in line with the broader move toward robust learning-based brain extraction that generalizes across contrasts and acquisitions, such as SynthStrip [32]. In this pipeline, the deep learning–based mask was combined with surface-based priors to improve robustness across pediatric ages and anatomies, mitigating both over-stripping of cortex and inclusion of non-brain tissue.

#### Step 7: CSF-based intensity normalization

Intensity normalization was performed in native space using cerebrospinal fluid (CSF) as the reference tissue. Whereas adult pipelines often normalize relative to white matter, white matter signal intensity varies substantially across childhood due to ongoing myelination. CSF T1 intensity is comparatively stable across development and therefore provides a more appropriate reference. For each subject, the mean CSF intensity was estimated from tissue masks and used to scale the entire image to a common reference level, preserving age-independent tissue contrast relationships.

#### Step 8: Quantitative quality control

Automated image quality metrics were computed using MRIQC [3]. MRIQC extracts a standardized set of image quality metrics (IQMs) from structural MRI, including measures of noise, sharpness, tissue contrast, and artifact burden. Metrics such as the coefficient of joint variation, FBER, and EFC were monitored to identify scans with potential preprocessing failures or acquisition artifacts. During pipeline development, these metrics were used as feedback signals to guide parameter tuning, ensuring that denoising and normalization improved signal quality without degrading anatomical contrast.

#### Step 9: Visual inspection

All preprocessed images additionally underwent expert visual inspection. Volumes were reviewed in multiple planes to identify artifacts or preprocessing failures not captured by automated metrics, with particular attention to cortical boundaries, inferior temporal and frontal regions, and ventricular spaces. Scans exhibiting severe motion artifacts, residual bias fields, or poor brain masking were excluded from template construction. This combined automated and manual quality control strategy reduces the risk that low-quality images degrade the final template.

#### Step 10: Re-centering and alignment

Finally, each image was recentered to a standardized coordinate origin to remove gross positional offsets across subjects. Re-centering reduces variability in registration initialization and improves convergence of subsequent nonlinear alignment during template construction.

Following preprocessing, we constructed age-specific and sex-specific pediatric brain templates using an iterative, groupwise diffeomorphic registration framework (Fig. 2). The goal of this stage was to generate unbiased population-average reference spaces that faithfully capture cohort-level anatomy while minimizing bias toward any individual subject.

**Fig. 2:**
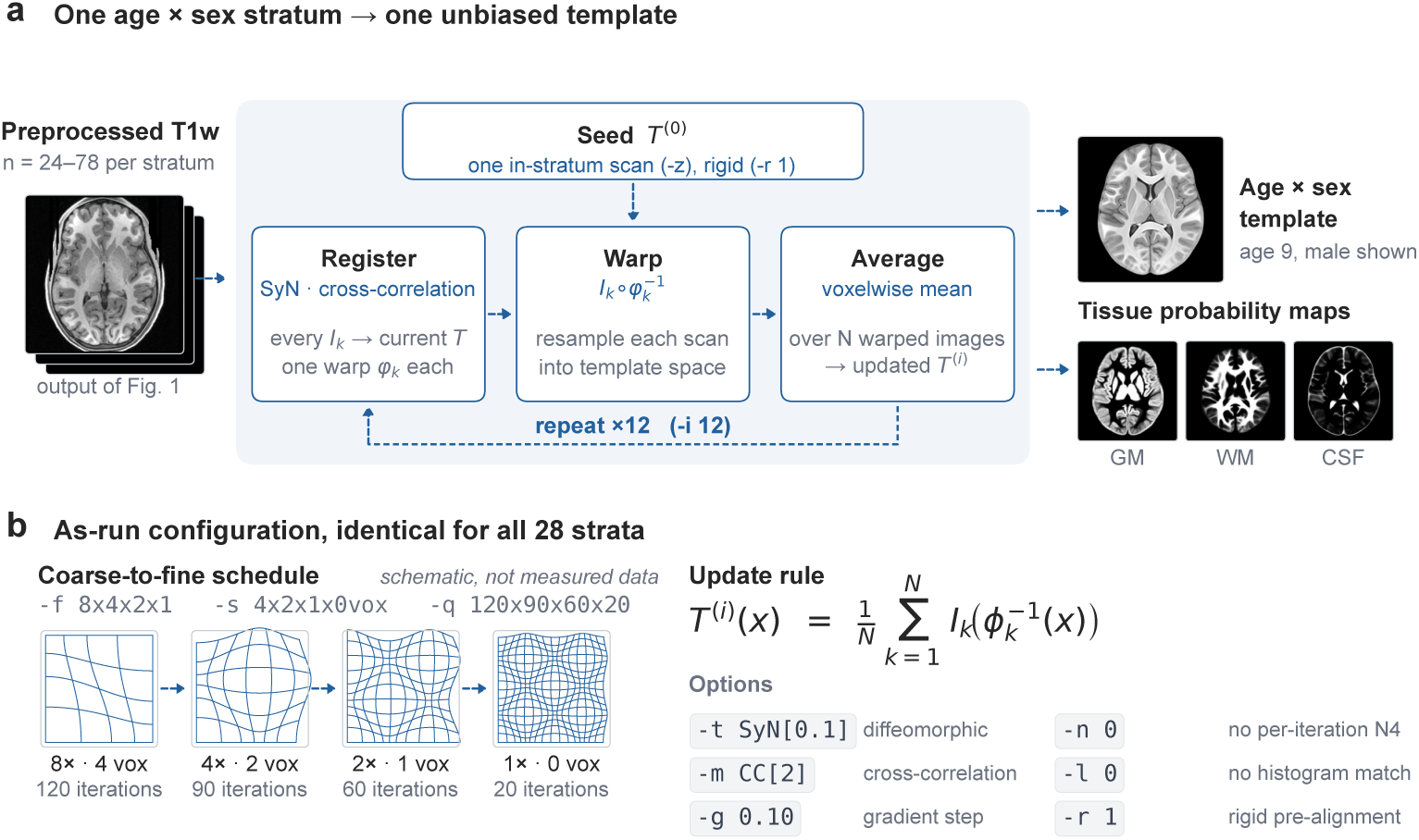
Iterative groupwise diffeomorphic template construction. **a**, Every preprocessed scan in one age × sex stratum is registered to the current template estimate, warped into template space and averaged; twelve such iterations converge on an unbiased cohort average. The seed is a single high-quality in-stratum scan rather than a blurred average, which accelerates convergence toward sharp consensus anatomy. Outputs are the template and its released tissue-probability maps; the age-9 male stratum is shown, at true physical scale. **b**, The as-run configuration, identical for all 28 published strata. The four grids illustrate what a coarse-to-fine schedule does and are schematic, not measured displacement fields. -n 0 disables per-iteration N4 rescaling so that the CSF-anchored contrast established in preprocessing (Fig. 1) survives averaging.

Template construction was performed using ANTs, building on the symmetric diffeomorphic normalization (SyN) framework introduced by Avants and colleagues for population-based brain image registration and atlas construction [5, 24]. In this approach, all subjects within a cohort are repeatedly aligned to a common template estimate, warped into template space, and averaged to update the template. Iterating this procedure yields an unbiased average that reflects the shared anatomical structure of the group rather than the geometry of an arbitrary reference.

We implemented this procedure using the ANTs multivariate template construction workflow (antsMultivariateTemplateConstruction2.sh), which has been widely used and validated for building population-specific brain templates across age groups and imaging modalities [2, 24]. While the core algorithmic framework follows established methods, we adapted the registration strategy to pediatric anatomy through cohort stratification and parameter tuning, as described in detail in the following section. We note that recent learning-based paradigms offer alternative routes to template construction and registration, including learned conditional (age/sex-conditioned) deformable templates such as AtlasMorph [33] and foundation models for medical image registration such as uniGradICON [34], alongside SynthMorph [35]; we nonetheless adopt regularized, controllable groupwise ANTs SyN here because it affords explicit, auditable control over regularization and intensity handling for curated pediatric template construction.

An overview of the iterative process—including subject-to-template registration, template updating, and convergence—is illustrated in Fig. 2. Registration parameterization, intensity-handling decisions, and pediatric-specific optimizations are presented in the subsequent section.

### 2.2 Controlled Denoising and Bias-Field Correction

All T1-weighted images underwent modest noise reduction and intensity inhomogeneity correction prior to further analysis. We applied a spatially adaptive non-local means (SANLM) filter [23] with minimal strength (filter parameter *v* = 1) to suppress high-frequency noise while preserving fine structures. This method adaptively modulates smoothing based on local noise estimates [23], thereby avoiding oversmoothing of developing sulci and thin cortical ribbons in children. Compared to standard non-local means, the adaptive variant better handles spatially varying noise (e.g. higher near coil elements) and was shown to improve SNR without blurring edges [23]. Next, we performed bias-field correction using the N4 algorithm [27] (ANTS 2.5 implementation), which iteratively estimates and divides out a smooth field of intensity non-uniformity. We used N4 with default B-spline fitting parameters (convergence to < 10^−6^ or 50 iterations per level) to remove intensity gradients caused by coil profile and facilitate consistent tissue intensities across the field of view. Notably, pediatric brains exhibit evolving tissue contrast with age (e.g. incomplete myelination yielding low white/gray contrast in infants), making them especially sensitive to bias-field distortions. By combining gentle SANLM denoising and N4 correction, we standardized images for downstream processing: noise was reduced enough to avoid spurious tissue segmentations, and residual bias was removed to not confound intensity-based measurements. These steps were intentionally conservative – the SANLM filter’s strength was throttled (one iteration at low variance) to avoid erasing subtle anatomical details, and N4’s stopping criteria were tuned to prevent overcorrection in very homogeneous infant brains. This controlled approach improved overall image quality while preserving biologically meaningful tissue-intensity differences critical for pediatric scans (e.g. distinguishing unmyelinated white matter from gray matter).

### 2.3 Explicit Background and CSF Modeling

#### Step 5: Background noise handling

Before template construction, we explicitly modeled the image background rather than leaving all outside-brain voxels at exactly zero. After skull stripping, each subject-specific brain mask was dilated by two voxels to define a narrow shell surrounding the brain. Voxels within this shell retain their original T1 intensities, and every voxel beyond it was set to a single small constant equal to 1% of the standard deviation of the background (air) intensities measured in the corresponding input image. This avoids a hard intensity discontinuity at the brain boundary and provides a realistic, non-zero noise floor, which can stabilize registration and intensity mapping when MRI intensities lack an absolute scale and when background is treated consistently across subjects [36].

#### Step 7: CSF-referenced intensity normalization

To bring all individual T1 images to a common intensity scale prior to averaging, we normalized each scan using cerebrospinal fluid (CSF) as a tissue landmark. An initial CSF estimate was obtained via intensity-based tissue classification (FSL FAST; [37]), then refined using FreeSurfer recon-all segmentation to isolate ventricular CSF labels in native space (FreeSurfer; [28, 38, 39]); modern contrast- and resolution-agnostic learning-based segmentation (e.g. SynthSeg [40]) offers an alternative route to such tissue labels. We computed the mean ventricular CSF intensity *µ*_CSF_ and scaled the entire image as *I*_norm_(*x*) = *I*(*x*)/µ_CSF_, yielding a dimensionless intensity scale anchored to an anatomically defined compartment. This tissue-referenced scaling reduces inter-subject variability due to scanner gain and acquisition differences and helps preserve consistent tissue contrast in the resulting cohort template. Related large-scale template efforts similarly applied global intensity scaling prior to averaging (e.g., ICBM/MNI resources) [41].

#### Role of FreeSurfer for segmentation, normalization, and QC

We emphasize that FreeSurfer was used primarily to provide anatomically grounded tissue/structure masks (via aseg.mgz) rather than to directly improve registration. In particular, the atlas-informed labeling enables a precise definition of ventricular CSF for intensity scaling and supports tissue-informed quality control. Using FreeSurfer-derived masks, we computed tissue-specific intensity summaries (e.g., within CSF and white matter) to flag outliers indicative of preprocessing failures (residual bias field, masking errors) before template construction [28, 38].

### 2.4 MRIQC-Guided Iterative Tuning and Triage

Rather than treating quality control (QC) as a final binary pass/fail assessment, we used automated QC as an *iterative diagnostic* to understand limitations of our initial preprocessing workflow and to guide revisions to both parameters *and* processing steps. We leveraged MRIQC (v0.16.1; [3]) to extract a rich set of image quality metrics (IQMs) from structural scans; recent benchmarking of automated T1-weighted QC methods supports the use of such IQM-based curation for research datasets [42]. These no-reference IQMs quantify complementary aspects of image quality [43], enabling a data-driven refinement of pediatric MRI preprocessing.

In particular, we focused on MRIQC-derived IQMs with direct interpretability for template fidelity—tissue contrast, intensity non-uniformity, and spatial clarity— and summarized representative pre- vs. post-processing ranges in Table 1a together with qualitative interpretation rules in Table 1b. We tracked contrast and intensity-distribution metrics such as the contrast-to-noise ratio (CNR), coefficient of joint variation (CJV), intensity non-uniformity (INU), and the white-matter to maximum intensity ratio (WM2max), alongside sharpness/artifact proxies such as the entropy focus criterion (EFC) and full-width at half-maximum (FWHM), and overall signal/background separation metrics such as the foreground-to-background energy ratio (FBER) and SNR [29, 43, 44]. Under standard MRIQC definitions, increases in CNR and WM2max (within expected ranges), together with decreases in CJV/INU and FWHM, are generally consistent with improved image quality [29].

**Table 1:** Representative MRIQC-derived image quality metrics before and after pre-processing, with qualitative interpretation. Arrows indicate whether higher (*↑*) or lower (*↓*) values are better under standard MRIQC definitions. **Several metrics are sensitive to background handling and are reported here for transparency; where preprocessing violates metric assumptions, post-processing values are interpreted cautiously (see text).**

| Metric ($\uparrow / \downarrow$ = better) | Range Pre | Range Post |
| --- | --- | --- |
| CNR ↑ | 0.91–1.95 | 0.91–2.34 |
| CJV ↓ | 0.55–0.98 | 0.47–0.89 |
| INU range ↓ | 0.25–0.83 | 0.03–0.07 |
| WM2max ↑ | 0.43–0.63 | 0.57–0.84 |
| FWHM avg ↓ | 3.64–4.44 mm | 3.11–4.27 mm |
| EFC ↓ | 0.47–0.54 | 0.78–0.82* |
| FBER ↑ | 2.3k–16.3k | invalid<sup>†</sup> |
| SNR ↑ | 4.56–6.83 | 1.66–3.02<sup>‡</sup> |
| tpm_overlap ↑ | 0.43–0.45 | 0.38–0.42 |

\*EFC increased post-processing, indicating that entropy-based sharpness metrics may be confounded by mask boundaries or background modeling; values are reported for transparency and not used as acceptance criteria.
<sup>†</sup>FBER became invalid post-processing due to nonstandard background intensity handling; this metric was excluded from threshold-based triage in the current version.
<sup>‡</sup>SNR decreased post-processing, consistent with conservative background noise reinjection and/or intensity operations; SNR was monitored but not optimized as a primary objective.

**Table 1:** Representative MRIQC-derived image quality metrics before and after preprocessing, with qualitative interpretation.
| Metric | $\Delta$ & interpretation |
| --- | --- |
| CNR | ↑ Tissue contrast improved, indicating effective bias-field correction and denoising. |
| CJV | ↓ Grey/white variance reduced, supporting more uniform intensity scaling. |
| INU | ↓ Consistent with successful N4 bias-field correction. |
| WM2max | ↑ White-matter peak intensity closer to the theoretical optimum. |
| FWHM avg | Small change; slight sharpening suggesting modest spatial smoothing. |
| EFC | ↑ Contrary to the expected direction; likely confounded by mask boundaries or background modeling. Reported for transparency and not used for acceptance in the current version. |
| FBER | Invalid after preprocessing due to altered background statistics; excluded from threshold-based triage in the current version. |
| SNR | ↓ Decreased post-processing; monitored as a diagnostic but not treated as a primary optimization target given background noise reinjection and intensity harmonization steps. |
| tpm_overlap | Slight drop, indicating registration or segmentation mismatch. |

We used these IQMs as quantitative feedback rather than as a single equation-driven optimization target. Specifically, we ran a representative subset of subjects through multiple candidate pipelines and compared their multi-metric MRIQC profiles to identify which configuration yielded the most desirable overall pattern across complementary IQMs. Table 1 illustrates this comparison for a representative subgroup (female, 9 years): the same subjects were processed with candidate workflows and their MRIQC outputs were examined to identify consistent improvements in contrast-and bias-related metrics (e.g., higher CNR, lower CJV/INU) without unacceptable degradation in sharpness proxies.

Crucially, this MRIQC-guided comparison informed *step-level* changes to the pipeline, not only parameter tuning. For example, trends in background- and mask-sensitive IQMs motivated explicit background padding/noise modeling (to avoid edge discontinuities), and contrast/inhomogeneity patterns motivated adding N4 bias-field correction as a dedicated step. After incorporating these changes, we re-ran the same representative subset to verify that the revised workflow improved the overall MRIQC profile in the intended directions.

#### Known limitations and planned refinements

Several MRIQC metrics assume background intensities are near zero and that mask boundaries do not introduce strong edge discontinuities. Because our pipeline explicitly models background statistics (Step 5) to preserve realistic noise properties for downstream processing, a subset of background- and entropy-sensitive IQMs (notably FBER and, in some cases, EFC) can become unstable or reflect preprocessing-induced distributional changes rather than acquisition quality. We therefore report these metrics for transparency (Table 1) but interpret them cautiously and do not treat them as primary acceptance criteria in the current version. Future refinements will (i) compute background-sensitive IQMs on a standardized background representation (e.g., using raw-background estimates prior to reinjection), and (ii) evaluate a composite QC index calibrated to expert ratings for pediatric anatomy.

Although we compared candidate pipelines using the multi-metric MRIQC profiles described above, we did not yet implement a single composite score for automated ranking or subject-level flagging. A natural next step is to aggregate multiple IQMs into a unified, interpretable index for prospective QC-driven optimization and automated outlier detection. In general form, one may define a composite score as

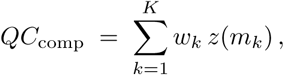

where *m_k_* denotes an IQM (e.g., CNR, CJV, EFC, FBER, SNR), *z*(*m_k_*) normalizes it to a common scale (e.g., *z*-scoring across subjects or relative to a reference distribution), and *w_k_* encodes importance and directionality (positive for metrics where higher values are better; negative for metrics where lower values are better). Such composite formulations can be calibrated against expert visual ratings [45] and would enable systematic pipeline selection as well as scan flagging in future work.

### 2.5 Demographic-Aware Deformation Cost Modeling

To investigate how demographic differences between template and subject affect registration difficulty, we quantified a deformation cost for each subject–template pairing and modeled it as a function of age and sex. We define deformation cost *D_i,j_* as the normalized energy or displacement required to warp subject *i* to template *j*. Both sides of that registration are gray-scale images: the moving image is the subject’s CSF-normalized, recentered T1 volume (subj_<id>_csfNorm_rc.nii.gz), the same intensity image used everywhere else in this work, and the fixed image is the corresponding template. Cost is accumulated within the template brain mask, and over the SyN warp field alone—the affine component, which carries a near-rigid coordinate-origin offset as well as global scale, is excluded from the displacement. We state the inputs explicitly because the metric is only interpretable when both images carry intensity texture: a cross-correlation metric has zero local variance inside a binary mask, so registering a brain *mask* to a gray-scale template would leave the optimizer with gradient information only at the brain boundary and the resulting displacement would reflect regularization rather than anatomy. In practice, we used two proxies for *D*: (1) the mean squared displacement of the deformation field (in mm^2^), normalized by brain volume, and (2) the log transform energy 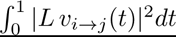 from the SyN optimization (reported by ANTs as the final metric value). Both measures gave consistent results, so we focus on the displacement metric for interpretability. For each subject *i*, we computed *D_i,j_* to both a matched template (same sex, closest age bin) and a mismatched template (opposite sex, same age bin), and examined the within-subject difference:

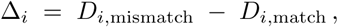

which isolates the cost penalty when using an opposite-sex template for the same individual. A positive Δ*_i_* indicates that a mismatched template yields higher deformation cost (worse fit) than a sex-appropriate template; in our data, Δ*_i_* was positive on average (see Results), consistent with prior reports that sex-specific templates can reduce registration burden [46].

To formally quantify demographic contributions while separating them from large subject-specific variability, we adopted a generative decomposition of *D_i,j_* into interpretable components:

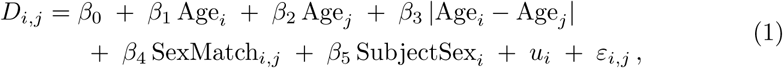

where fixed effects quantify the average influence of subject age, template age, age mismatch, and sex matching, and the subject-specific random intercept *u_i_* captures baseline differences in registration difficulty. Conceptually, this framework treats deformation cost as the sum of subject anatomy, age-dependent maturation, template age fitness, an age mismatch penalty, a sex matching effect, and noise. We fit Eq. 1 by restricted maximum likelihood on the masked-mean SyN displacement metric (*D_i,j_* = mean displacement in mm; never the composite affine◦SyN cost, which adds a subject-specific term largely unrelated to template fit—see Supplementary Fig. S2) over the sex-specific pediatric template subset, with the deformation cost *D_i,j_* on the response side; the fitted coefficients are reported in the Results (Sec. 3.5). The sex-neutral NKI reference templates, for which SexMatch and template sex are undefined, were excluded from this primary fit so that the SexMatch term does not confound template source with sex matching.

Importantly, this demographic-aware deformation modeling was applied exclusively at the evaluation stage (after template construction): it serves as a diagnostic tool to assess template suitability and potential demographic bias, rather than an optimization objective that alters registration itself.

### 2.6 Jacobian-Based Regularity QC

To verify the topological correctness and stability of the deformations, we analyzed the Jacobian determinant maps of all nonlinear warp fields generated during template construction. The Jacobian determinant *J* (*x*) at each voxel *x* indicates local volume change (expansion if *J >* 1, contraction if *J <* 1) introduced by the transform. We examined three summary measures: (1) the mean log-Jacobian, averaged over brain voxels within each warp, (2) the within-warp standard deviation of log *J*, and (3) the percentage of voxels with non-diffeomorphic transforms (i.e. with *J ≤* 0, which would imply folding). These were computed over all 1,272 subject-to-template construction warps spanning the 28 age × sex strata (Fig. 3), from the SyN warp field alone, with the affine component excluded. That convention matters when comparing Jacobians across analyses: the affine carries global scale and shear, so a SyN-only and a composed affine ◦ SyN Jacobian are different quantities rather than rescalings of one another, and the two cannot be compared directly. For a strictly volume-preserving registration one would expect mean log *J ≈* 0. The observed mean was −0.032, corresponding to a small (∼3%) systematic contraction rather than an unbiased mapping; this reflects mild global compression arising from subject–template size differences. The effect was uniform across the library (per-stratum means −0.020 to −0.037, the extreme being the rebuilt age-11 female stratum) and showed no sex dependence (male −0.032, *n* = 757; female −0.032, *n* = 515), so it does not differentially distort either sex-specific arm of the library. The within-warp standard deviation of log *J*, which reflects the magnitude of local volume-change variability, averaged 0.276 across warps (per-warp range 0.228–0.339; per-stratum means 0.254–0.288). This magnitude is consistent with the greater shape variability of pediatric anatomy and denotes a smooth, plausible deformation field. Crucially, the fraction of non-diffeomorphic voxels was exactly 0% in every one of the 1,272 warps—no instance of *J ≤* 0 anywhere in the library. The diffeomorphic SyN algorithm theoretically guarantees invertible transforms [5], but numerical issues can occasionally produce tiny negative Jacobians. Every warp used to build the templates was therefore a one-to-one smooth mapping. These findings validate that the registration procedure produced well-behaved deformations without introducing artifacts, a critical requirement for subsequent analysis of deformation-based morphometrics.

**Fig. 3:**
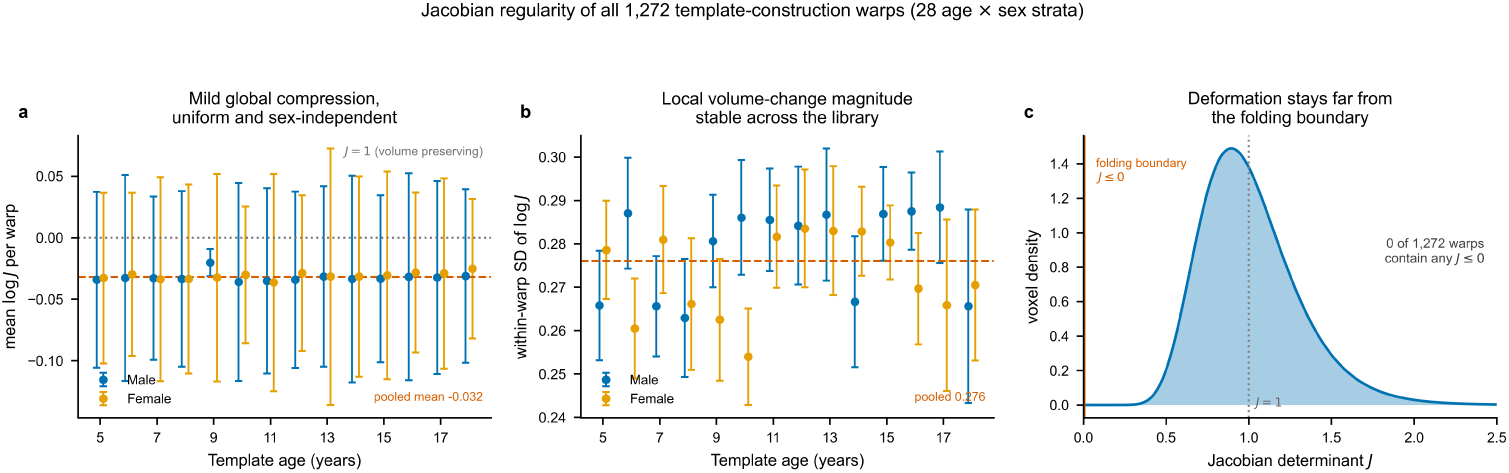
Jacobian regularity of every warp used to build the library. Computed over all 1,272 subject-to-template construction warps across the 28 age × sex strata. **(a)** Per-stratum mean log *J*; points are stratum means and bars the between-warp SD. The deformation is a mild global compression (pooled mean −0.032, dashed line) that is uniform across ages and effectively identical between sexes (male −0.032, *n* = 757; female −0.032, *n* = 515), rather than the volume-preserving mapping an unbiased registration would give (*J* = 1, dotted line). The age-9 male stratum is built from its 92 reconstructed (_rc) warps rather than the full 129 forward warps, which is why its between-warp spread is tighter. **(b)** Within-warp SD of log *J*, the magnitude of local volume change, is stable across the library (pooled 0.276; per-stratum means 0.254–0.288). Excluding the age-9 male stratum, the pooled mean log *J* is −0.033 rather than −0.032, so the differently constructed warp set does not drive the result. **(c)** Pooled voxelwise distribution of the Jacobian determinant, as a voxel-count-weighted mixture of the per-warp log-normals. The mass sits near *J* = 1 and stays far from the folding boundary: no voxel in any of the 1,272 warps had *J ≤* 0.

### 2.7 Reproducibility and computing environment

We placed strong emphasis on computational reproducibility and scalability across heterogeneous systems. As illustrated in Fig. 4 (workflow schematic), the entire pipeline was implemented in a fully containerized form. We developed and validated the processing environment locally using Docker, selecting a Linux base image (Ubuntu 22.04) to maximize compatibility across neuroimaging tools and to avoid host-specific dependency drift. The Docker image bundled all required software, including ANTs (v2.5.0), FreeSurfer (v7.4.1), FSL (v6.0.7), AFNI (v23.2.04), SPM12 with CAT12 (v12.9), and custom Python scripts for preprocessing, quality control, and orchestration. The CAT12 spatially adaptive non-local-means (SANLM) denoiser (preprocessing Step 2) runs under MATLAB R2023b in the as-run image; to remove this proprietary dependency for reuse, we additionally provide a license-free build in which SANLM is instead performed by ANTs DenoiseImage, which implements the same adaptive non-local-means algorithm.

**Fig. 4:**
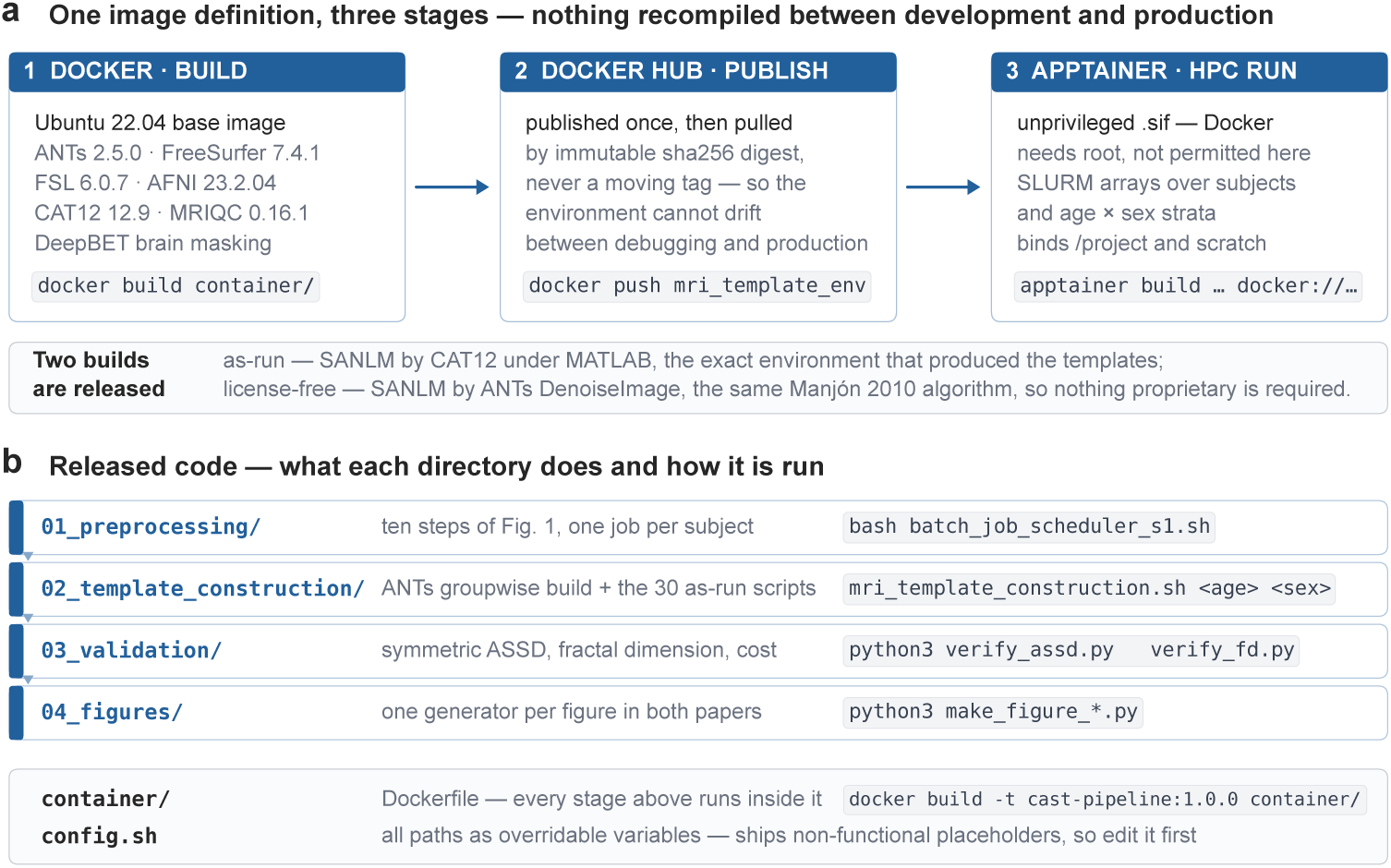
One environment across development and production, and how the released code is run. **a**, The processing environment is built once as a Docker image on an Ubuntu 22.04 base bundling the full neuroimaging stack at the versions shown, published to Docker Hub and pulled by immutable sha256 digest rather than by a moving tag, then converted into an Apptainer/Singularity image (.sif) for unprivileged execution on the shared HPC cluster, where Docker is not permitted because it requires root. Major pipeline stages run as SLURM batch jobs and job arrays, parallel across subjects and age/sex strata, with the project filesystem and per-job scratch bound at runtime. Two images are released: the as-run build, in which SANLM denoising runs under MATLAB, and a licence-free build in which the same algorithm is supplied by ANTs DenoiseImage. **b**, Layout of the released pipeline repository with the entry point of each stage. All stages execute inside the container of **a**, and config.sh ships non-functional path placeholders that must be set for the local environment.

To enable execution on shared high-performance computing (HPC) infrastructure where Docker is not permitted, the Docker image was published to Docker Hub and converted into an Apptainer (Singularity) container (.sif). This strategy ensured byte-identical runtime environments between local development, debugging, and large-scale cluster execution, eliminating discrepancies due to operating system, library versions, or compiler differences. Where possible, we verified that the container image digest/hash was identical across environments to emphasize determinism. Apptainer containers were executed without elevated privileges on the cluster, with project directories and temporary scratch space bound at runtime.

All large-scale processing was performed on the University of Houston HPC system operated by the Research Computing Data Core. Jobs were managed via the SLURM scheduler, with major pipeline stages (preprocessing, MRIQC-based quality control, FreeSurfer segmentation, and ANTs-based registration and template construction) submitted as independent batch jobs or job arrays to enable efficient parallelization across subjects and age/sex strata. This containerized, scheduler-driven design follows the same reproducibility and provenance-tracking principles advocated for large-scale BIDS-App neuroimaging workflows, which have been demonstrated at scale on the full HBN cohort (*n* = 2,565) [47]. Resource requests varied by stage; the SLURM directives actually used are provided verbatim with the released pipeline code (see Code Availability). Cluster accounting over the period retained by the scheduler (from 2024-11-09 onward) records approximately 5.8 × 10^5^ CPU-hours across roughly 7.1 × 10^4^ jobs for this user account. That figure spans all of this account’s activity rather than this project alone, and predates the retention window only in part, so it should be read as an order-of-magnitude indication of compute scale rather than a project-specific total. Data and intermediate derivatives were stored on the shared project filesystem (/project/contreras-vidal); the project occupied ∼15 TB of a 20 TB allocation at the time of analysis, reflecting retention of raw MRI data, intermediate preprocessing outputs, deformation fields, and multiple template iterations to support reproducibility and auditing.

### 2.8 Limitations

While our framework addresses many pediatric MRI challenges, we acknowledge certain limitations. First, the age range of our templates (5–18 years) does not cover infancy; applying our methods to neonates may require retuning of parameters (e.g. neonates have radically different tissue contrasts that might need different CSF normalization or atlas priors). We also did not incorporate longitudinal consistency explicitly — for studies with serial scans of the same child, incorporating longitudinal registration constraints could further improve anatomical alignment across time (our pipeline treats each timepoint independently). Additionally, our QC composite score weighting was empirically determined on a subset and might not generalize to data from different scanners or populations; others may need to recalibrate the weights *w_k_* for their specific dataset. In terms of segmentation, FreeSurfer — although usable with QC — is known to have reduced accuracy in very young children (under ∼4–5 years) [48]; our reliance on FAST for those cases means we forego surface parcellation detail in the youngest cohort. A related point is that we lacked ground-truth segmentations for validation, so our Dice metrics are relative to template priors, not absolute truth — true accuracy could be lower if the priors are imperfect. The deformation-based analyses (age/sex effects) assume that registration cost is a meaningful proxy for anatomical differences; this is generally valid, but can be confounded by scanner or processing artifacts. We attempted to mitigate this by rigorous QC and using robust metrics, but subtle biases might remain; for instance, a systematic sex difference in image quality could in principle influence the sex-matching coefficient η, although we saw no evidence of this. Another limitation is computational: the high-dimensional registration (SyN) is time-consuming. Although we consider the accuracy worth the cost, some settings (like the very large number of iterations we used) could potentially be trimmed for speed at minimal accuracy loss; we did not exhaustively explore that tradeoff. The decision to add synthetic noise in the background, while improving registration, could conceivably introduce bias in intensity-based analyses outside the brain (though we do not analyze outside brain). We assume the background noise is truly uninformative; to avoid introducing a spurious offset we used low-amplitude noise with a small positive mean matched to the measured air-background statistics of each input image (Step 5), kept non-negative so that it remains physically plausible under integer intensity storage rather than producing negative values. Finally, our templates, being averaged, blur fine structural details (especially in cortex where developmental variability is high). This is an inherent limitation of template building – while we preserved major anatomy well, certain high-frequency features (e.g. sharp gyral patterns unique to individuals) will be smoothed out. Researchers interested in those might require personalized templates or multi-atlas approaches. Despite these limitations, we believe our framework provides a solid, reproducible foundation for pediatric MRI processing. We explicitly invite the community to utilize and modify our pipeline (via the provided code) to suit other pediatric sub-populations, and we foresee future improvements such as integrating advanced denoising (e.g. deep-learning methods) or more dynamic atlas frameworks (4D atlases that continuously model age [26]). All analyses in this study should be interpreted in light of these considerations – for example, the sex differences found are template-fit differences, not definitive biological statements. By being transparent about limitations and assumptions, we aim to ensure that the methods and findings are used appropriately and spur further methodological development in pediatric neuroimaging.

## 3 Results

### 3.1 Dataset and template library

Templates were constructed using structural MRI data from the Healthy Brain Network (HBN), a large-scale, community-based pediatric neuroimaging initiative led by the Child Mind Institute and distributed through the International Neuroimaging Data-Sharing Initiative (INDI) [1]. The HBN dataset was designed to support transdiagnostic research in pediatric mental health and learning disorders and includes multimodal data spanning structural MRI, EEG, cognitive assessments, and extensive phenotypic and behavioral measures [1].

#### MRI acquisition

All contributing T1-weighted scans were acquired on 3 T Siemens MRI systems at three Healthy Brain Network sites—the Citigroup Cornell Brain Imaging Center (CBIC), the City University of New York Advanced Science Research Center (CUNY), and the Rutgers University Brain Imaging Center (RUBIC). Data collected on the 1.5 T mobile scanner at the Staten Island site were excluded, so that the cohort comprises 3 T acquisitions only. Native T1-weighted voxel sizes ranged from 0.8 to 1.0 mm isotropic across the contributing 3 T acquisitions, with full brain coverage (176–320 slices) and MPRAGE-style contrast optimized for pediatric anatomy (TR ≈ 2500–2730 ms, TI ≈ 1000–1060 ms, flip angle 7–8^◦^; multiband and partial Fourier disabled). Each age- and sex-specific template was constructed and released on its native acquisition grid rather than being resampled to a common atlas space: templates are distributed at either 0.8 mm isotropic resolution (224 × 320 × 213 voxels) or 1.0 mm isotropic resolution (176×256×170 voxels), depending on the underlying acquisition, in NIfTI format in approximate alignment with MNI orientation but not resampled to the 1 mm MNI152 182 × 218 × 182 grid, with file organization and metadata following community standards for neuroimaging data [49]. These parameters are consistent with the publicly released HBN MRI protocol [50].

#### Inclusion criteria

Participants eligible for inclusion were male or female children and adolescents aged 5–21 years at enrollment [1]. All participants (or their legal guardians) provided informed consent, with age-appropriate assent obtained from minors. Participants were required to be fluent in English, with accommodations available for Spanish-speaking parents when appropriate [1]. For the present study, only children free of gross structural abnormality within the 5–18 year age range were considered for template construction, following quality control and anatomical screening. HBN is a community sample enriched for mental-health and learning concerns, so this screening establishes the absence of gross structural abnormality rather than a strictly healthy or clinically vetted population.

#### Quality control and cohort refinement

All available T1-weighted images within the target age range underwent automated quality assessment followed by expert visual inspection (see Methods). Images exhibiting excessive motion artifacts, incomplete brain coverage, preprocessing failures, or atypical anatomy were excluded prior to template construction. This two-stage screening process ensured that the final templates reflected high-fidelity pediatric anatomy rather than artifacts introduced by marginal-quality scans.

#### Final age- and sex-specific template cohorts

After quality control, subjects were stratified into one-year age bins and further subdivided by sex. Table 2 summarizes the final number of subjects contributing to each template (1,272 in total), which formed the basis for all subsequent analyses. Cohort sizes were largest between ages 7 and 11 years (124–168 subjects per year) and gradually declined in late adolescence, reflecting the underlying age distribution of the HBN dataset.

**Table 2:**
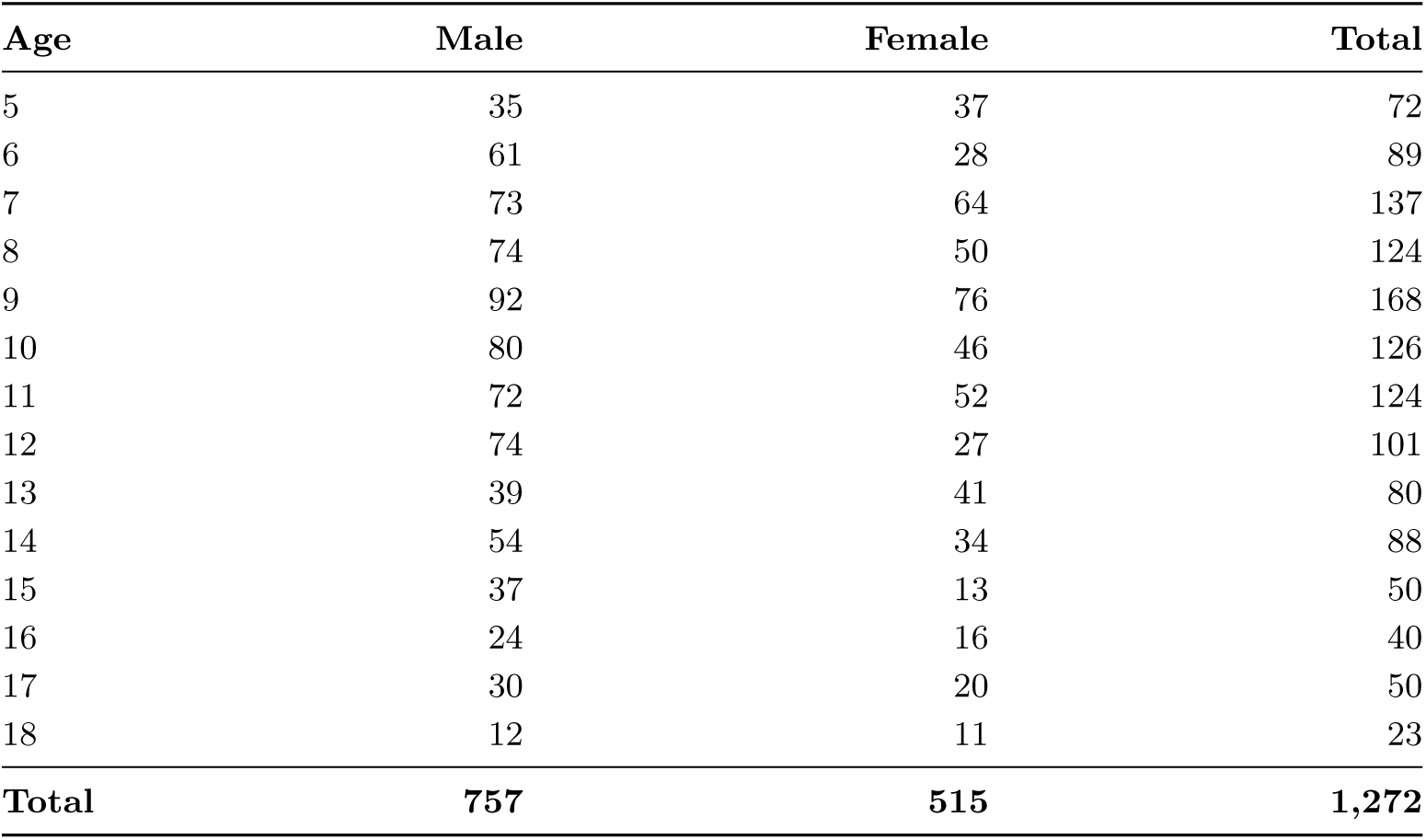
Final subjects used for age- and sex-specific template construction.

| Age | Male | Female | Total |
| --- | --- | --- | --- |
| 5 | 35 | 37 | 72 |
| 6 | 61 | 28 | 89 |
| 7 | 73 | 64 | 137 |
| 8 | 74 | 50 | 124 |
| 9 | 92 | 76 | 168 |
| 10 | 80 | 46 | 126 |
| 11 | 72 | 52 | 124 |
| 12 | 74 | 27 | 101 |
| 13 | 39 | 41 | 80 |
| 14 | 54 | 34 | 88 |
| 15 | 37 | 13 | 50 |
| 16 | 24 | 16 | 40 |
| 17 | 30 | 20 | 50 |
| 18 | 12 | 11 | 23 |
| **Total** | **757** | **515** | **1,272** |

Using the Healthy Brain Network (HBN) dataset [1] described above, we generated age- and sex-specific templates for each one-year bin from 5 through 18 years from a candidate pool of over 2,400 T1-weighted scans, of which 1,272 passed preprocessing and quality triage and contributed to the templates. The complete age-by-sex template series—displayed at a matched axial level in true physical (millimetre) size, together with the corresponding gray-matter, white-matter, and CSF tissue-probability maps for every template—is presented in the companion data descriptor (Hu & Contreras-Vidal, in review, *Scientific Data*), which owns the at-a-glance library montage and tissue-map figures.

### 3.2 Templates encode normative development and sex dimorphism

Beyond visual fidelity, the templates reproduce the canonical anatomical trajectory of school-age and adolescent development. Tissue-volume morphometry of the 28 templates (FSL FAST) shows white-matter volume increasing with template age (*r* = +0.68), gray-matter volume declining (*r* = −0.54), intracranial volume growing (*r* = +0.51), and the white-to-gray volume ratio—a proxy for ongoing myelination—rising steadily (*r* = +0.82). These template-derived trajectories track the independently measured subject growth norms (FreeSurfer, *n* = 1,473; shown in the companion data descriptor [Hu & Contreras-Vidal, in review, *Scientific Data*]), confirming that each template encodes age-appropriate anatomy rather than a generic average. Structural complexity, quantified by the fractal dimension of the white-matter surface, likewise increases with age (*r* = +0.70) and exceeds that of the age-matched NKI reference. Scoring both libraries with the identical multi-Otsu white-matter generator, BRAIN CAST has the higher fractal dimension at every one of the eleven ages spanned by both libraries (paired mean difference +0.041, 11/11 ages; paired *p* = 4 × 10^−5^), an advantage that widens on a common 1.0 mm grid (+0.058, 11/11 ages, *p* = 1 × 10^−6^) and is therefore not a voxel-size artifact. This indicates that the fine-grained age stratification preserves developmental folding detail that broader-age references blur. The templates also capture established sex dimorphism: male templates are larger than female templates at essentially every age (intracranial volume +9.7%, gray matter +10.4%, white matter +10.0% on average; Fig. 5), consistent with reported sex differences in pediatric brain size [13, 14]. This sex-specific structural detail is, by construction, unavailable from a single sex-neutral reference. Together these morphometric trends establish that the library is genuinely age- and sex-specific—a prerequisite for the representativeness and validity analyses that follow. By encoding age- and sex-resolved normative anatomy, these references are well suited to downstream normative-modeling and individual-deviation analyses [51], complementing empirically benchmarked sex-specific normative-modeling resources [52].

**Fig. 5:**
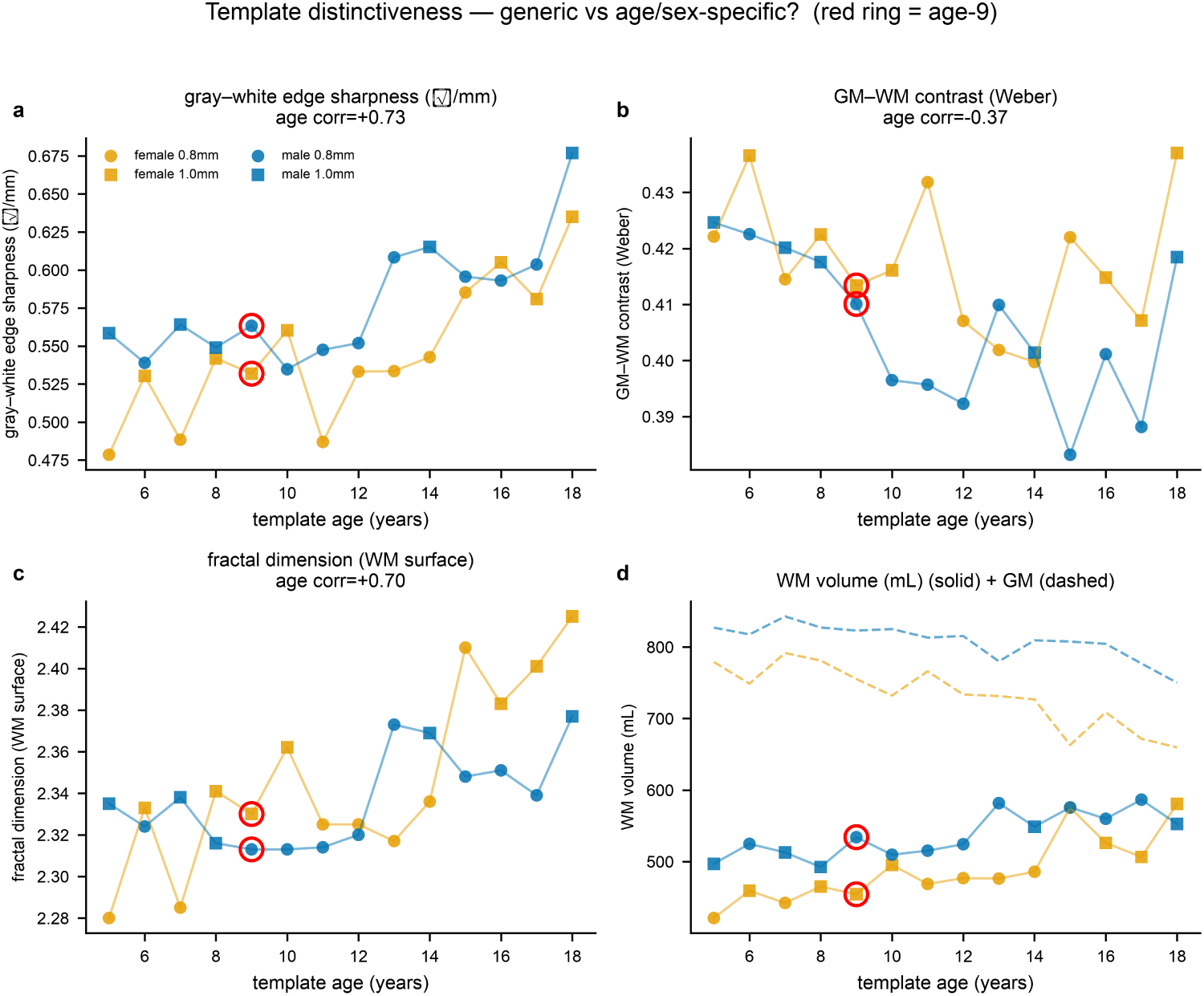
Templates are age- and sex-distinct, not generic. Per-template gray–white edge sharpness, GM–WM Weber contrast, white-matter fractal dimension, and WM/GM tissue volumes across all 28 strata (ages 5–18), separated by sex. Age-correlations confirm developmentally faithful trajectories; the age-9 template (red ring) lies within the scatter of its age-7–11 neighbors, i.e. it is not an outlier. Marker shape encodes template voxel resolution (0.8 vs 1.0 mm). Edge sharpness is the robust [0, 1]-normalised gradient on a common 1.0 mm grid and fractal dimension uses multi-Otsu segmentation—the same definitions as the companion data descriptor, so the two papers’ morphometry figures are directly comparable; these are not the raw-gradient and FSL-FAST variants, whose numerical ranges differ.

#### Qualitative comparison at age nine

Qualitatively, the age-nine female and male templates (built from 76 and 92 participants, respectively) show crisp cortical and subcortical boundaries and tissue contrast comparable to the age-nine NKI reference; a three-plane visual comparison is provided in the companion data descriptor (Hu & Contreras-Vidal, in review, *Scientific Data*), while the quantitative head-to-head below carries the comparative evidence.

### 3.3 Representativeness: sub-voxel gray–white fidelity

A template’s practical value is the structural bias it leaves after a subject is normalized to it—not its apparent sharpness, which can be high even for an over-smoothed or mis-shapen average (the “reliable but wrong” failure mode). We therefore measured downstream structural fidelity directly. For each of 271 held-out subjects (ages 5–12, both sexes), we reconstructed the cortical gray–white surface with FreeSurfer, brought it into the age- and sex-matched template via the same diffeomorphic registration used throughout, and computed the distance between the template’s gray–white boundary and the warped subject surface, restricted to the cortical interface (Fig. 6). Table 3 reconciles the held-out samples used across the validation analyses that follow.

**Fig. 6:**
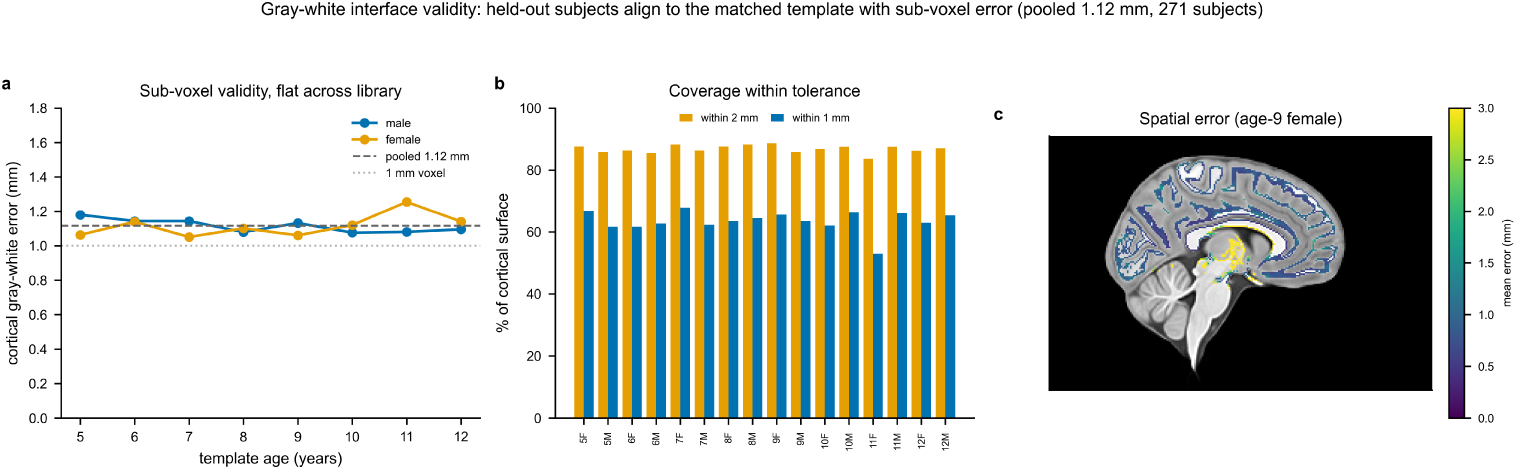
Gray–white interface validity (centerpiece). 271 held-out subjects (ages 5–12) aligned to the matched template. **a**, Per-stratum mean cortical error versus age: sub-voxel (≈1.1 mm) and flat across the library, apart from the rebuilt age-11 female template (1.26 mm, *n* = 10 held-out subjects). **b**, Percentage of the cortical surface within 1 and 2 mm per stratum. **c**, Example spatial mean-error map (age-9 female template, sagittal slice).

**Table 3:** Analysis cohorts. Held-out samples used across the validation analyses, all drawn from the HBN Release-11 held-out pool; the template-construction and growth-norm cohorts are separate.

| Analysis | N | Age range | Notes |
| --- | --- | --- | --- |
| Template construction | 1,272 (of 2,444) | 5–18 | the released library |
| Developmental growth norms (FreeSurfer) | 1,473 | 5–18 | separate FreeSurfer cohort, larger than the 1,272 contributing scans |
| Held-out validation pool (HBN R11) | 458 | 5–22 | superset of the rows below |
| Gray–white interface validity | 271 | 5–12 | both sexes |
| Sex × subject–sex interaction | 271 (99 F, 172 M) | 5–12 | cross-sex comparison |
| BRAIN CAST vs NKI head-to-head | 144 paired | 5–10 | grid-matched 0.8 mm |
| BRAIN CAST vs Fonov head-to-head | 209 | 5–12 | — |
| Tissue-specific Jacobian | 81 | 5–12 | — |
| Deformation-cost mixed model | 3,190 regs / 319 subjects | 5–12 | subjects × template ages |
| SyN-saturation pilot | 33 regs | — | translation-artifact check |

Across all 16 age–sex strata, the mean cortical interface error was **1.12 mm** (median 0.77 mm), with 84–89% of the cortical surface within 2 mm and 53–68% within 1 mm—sub-voxel fidelity at the 0.8–1.0 mm template resolution. The error was uniform across fifteen of the sixteen strata (per-stratum mean 1.05–1.18 mm), indicating consistent representativeness at every age and sex. The exception is the rebuilt age-11 female template at 1.26 mm (53% within 1 mm); it carries the joint-smallest evaluation set in the sweep (*n* = 10 held-out subjects), so the estimate is the noisiest in the library and is reported as measured. The signed error was small, spanning −0.26 to +0.04 mm: the template gray–white boundary lies predominantly interior to the subject’s (one stratum marginally exterior), a minor, spatially diffuse bias.

This metric also localizes *where*sex matters. Registering each subject to the matched-sex and the opposite-sex template (affine sweep, isolating shape from size), the signal is a sex × subject-sex *interaction* (linear mixed-effects model with a per-subject random intercept, *p* = 7*imes*10^−3^)—not a uniform matched-sex advantage. Female subjects fit the female template significantly better (by 0.049 mm cortical, 0.129 mm full; Wilcoxon *p* = 1*imes*10^−17^, better in 95/99 subjects), whereas male subjects show a *reversal*, fitting the female (opposite-sex) template ≈ 0.038 mm better (*p* = 2*imes*10^−28^; Fig. 8). The reversal arises because the female template is an easier registration target for everyone (by ≈ 0.042 mm), a confound we disclose explicitly: the robust evidence for sex-specific construction is that female cortical shape is fit measurably better by its own template, whereas sex-matching does not benefit males on this metric. A single sex-neutral reference, such as NKI, structurally cannot provide sex specificity at all. This is also the domain in which sex effects are demonstrable at a magnitude that matters: the deformation-cost analysis below resolves a sex-matching term of only 0.014 mm, under a fifth of the structural effect and under a fiftieth of a voxel, because the diffeomorphic warp largely equalizes it.

Three caveats bound the interpretation. First, the subject surface is brought into template space with the same registration whose target is the template, so the residual reflects template–subject correspondence after best alignment; because the registration method is held fixed, comparisons across template age and sex remain valid. Second, the template-side boundary is a FAST segmentation (0.8–1.0 mm voxels) whereas the subject side is a continuous surface, so the reported distance is from template boundary voxels to the subject’s true interface. Third, the templates mix 0.8 and 1.0 mm resolution; the metric is in physical millimetres and is therefore largely robust to this.

### 3.4 Head-to-head against existing references

To place BRAIN CAST on the same footing as widely used references, we ran a grid-matched head-to-head benchmark on a separate set of held-out HBN subjects (resampled to a common 0.8 mm grid; *n* = 144 paired, ages 5–10), comparing the gray–white interface error against the contemporary single-template NKI reference and the broad-age Fonov/NIHPD reference (Table 4; lower is better). The BRAIN CAST–NKI comparison depends critically on the *direction* in which the interface distance is measured. Measured from the template boundary to the subject surface—the conventional one-sided form—NKI is closer by +0.049 mm on the full interface and +0.024 mm restricted to cortex (roughly 3–6% of a 0.8 mm voxel). But this one-sided distance structurally penalizes the template carrying the larger, more finely articulated cortical boundary: BRAIN CAST’s boundary sits marginally interior to the subjects’ (median signed offset −0.13 mm; the bias is small and spatially diffuse, Fig. 7) and carries more sulcal detail, so more of its boundary points lie far from the nearest subject vertex. Measured in the opposite direction (subject surface to template boundary) the ranking *inverts*—BRAIN CAST is closer by 0.035 mm cortical, in 128/144 subjects. The direction-symmetric Average Symmetric Surface Distance (ASSD), which averages both, is a statistical tie: BRAIN CAST−NKI = −0.005 mm by the mean (BRAIN CAST lower in 85/144) and +0.011 mm by the median (NKI lower in 105/144)—both under 1.5% of a voxel, with the sign determined only by the summary statistic— and no age stratum favours NKI (Fig. 9). We therefore read BRAIN CAST and NKI as *equivalent* in representativeness on this metric, the apparent one-sided NKI edge being an artifact of measurement direction rather than a difference in fit. Against the broad-age Fonov/NIHPD reference the result is decisive and direction-independent: BRAIN CAST is better by 0.150 mm (cortical) and in all 209/209 subjects. Against the nearest age-specific competitor—the Śanchez/Richards Neurodevelopmental MRI Database [2]—the same pattern resolves in BRAIN CAST’s favour. On the overlap of the two libraries’ coverage (*n* = 189 held-out subjects, ages 5–10 — the range over which the Neurodevelopmental MRI Database reference templates used here overlap the held-out cohort), the one-directional cortical median is again a near-tie (BRAIN CAST 0.762 vs Śanchez 0.756 mm), but the direction-symmetric ASSD favours BRAIN CAST *decisively* —1.099 vs 1.389 mm, a 0.29 mm advantage (95% CI 0.28–0.30) in all 189/189 subjects—because Śanchez’s reverse subject→template error is far larger (1.75 vs 1.10 mm). On the fair symmetric metric BRAIN CAST thus ties the best single-template reference (NKI) and decisively beats both the nearest age-specific competitor (Śanchez) and the broad-age reference (Fonov): the symmetric-ASSD ranking places BRAIN CAST and NKI together ahead of Śanchez, with Fonov last.

**Fig. 7:**
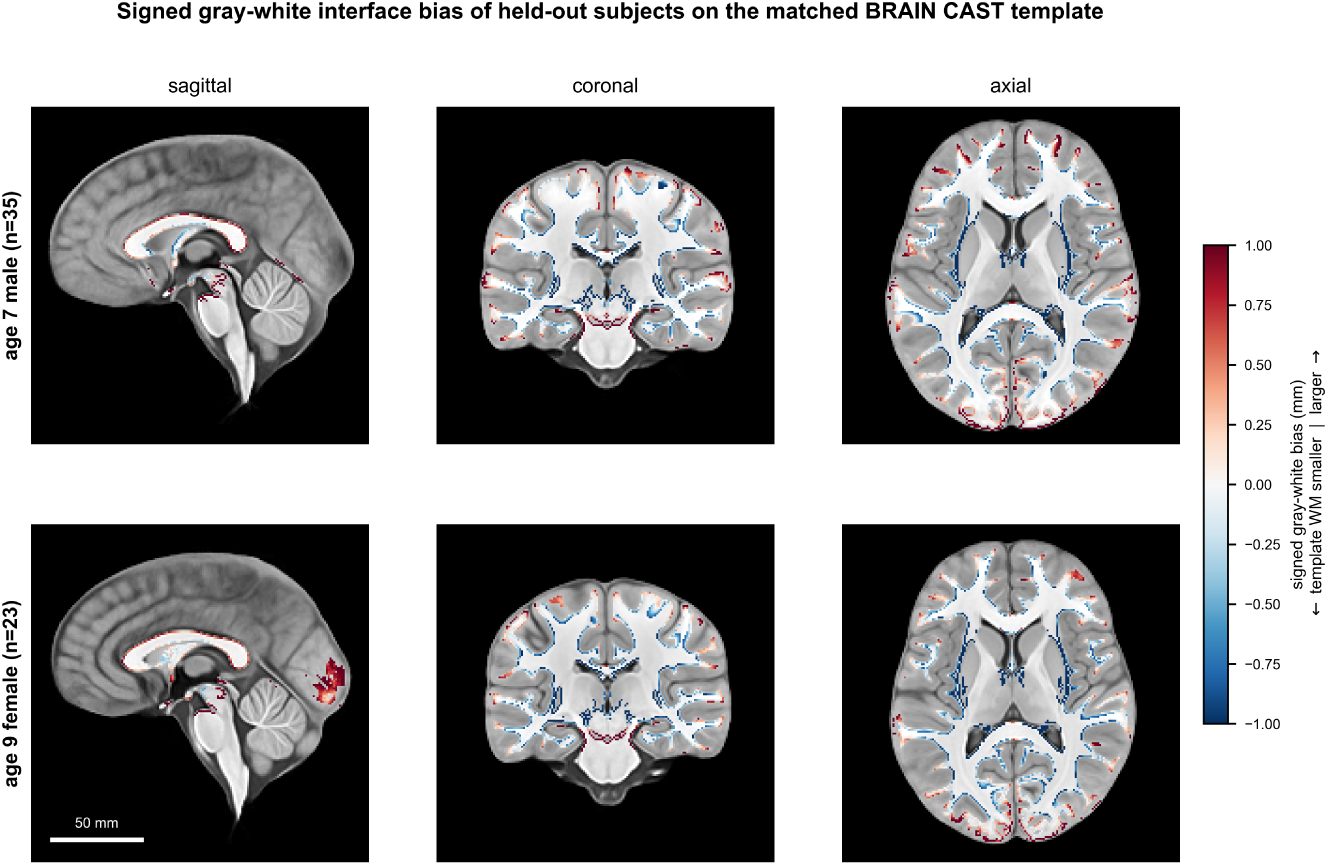
Signed gray–white interface bias is small and spatially diffuse. Per-voxel mean *signed* offset between the held-out subjects’ white surface and the matched BRAIN CAST template boundary (red, template WM larger; blue, smaller), for two example strata (age-7 male, *n* = 35; age-9 female, *n* = 23). The bias is sub-voxel along the cortical ribbon with no large systematic regions, and is mildly negative on average—the template boundary sits just inside the subjects’ own (cortical mean signed offset −0.08 and −0.26 mm for the two strata shown; +0.03 to −0.26 mm across all sixteen). This complements the magnitude map in Fig. 6: low residual structural bias, not merely small unsigned error.

**Fig. 8:**
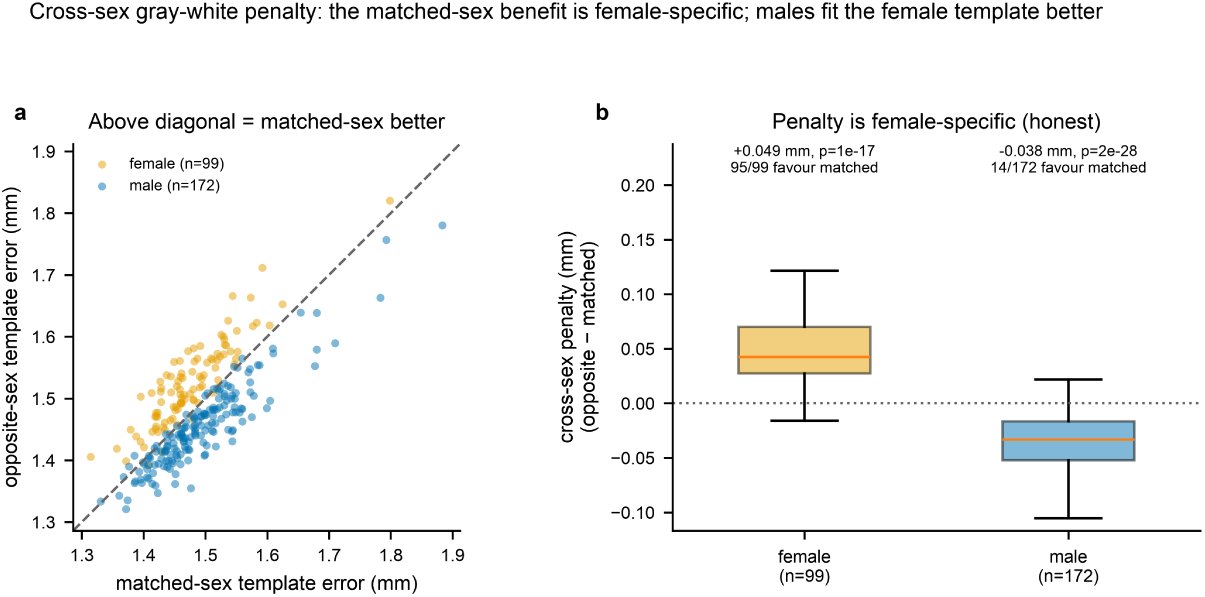
Cross-sex gray–white interaction. Cortical interface error to the matched-sex versus opposite-sex template (affine sweep, 271 held-out subjects). The signal is a sex × subject-sex interaction (mixed-effects *p* = 7 × 10^−3^), not a uniform matched-sex advantage. **a**, Paired per-subject errors; points above the diagonal are fit better by the matched-sex template—female subjects (orange) lie predominantly above it, male subjects (blue) below. **b**, The cross-sex penalty (opposite − matched) by subject sex: female subjects fit the female template better by 0.049 mm cortical (Wilcoxon *p* = 1 × 10^−17^, 95/99 subjects), whereas male subjects show a reversal, fitting the female template 0.038 mm better (*p* = 2 × 10^−28^, only 14/172 favouring the matched template), because the female template is an easier target for everyone (by 0.042 mm; disclosed confound). A sex-neutral reference cannot provide sex specificity at all.

**Fig. 9:**
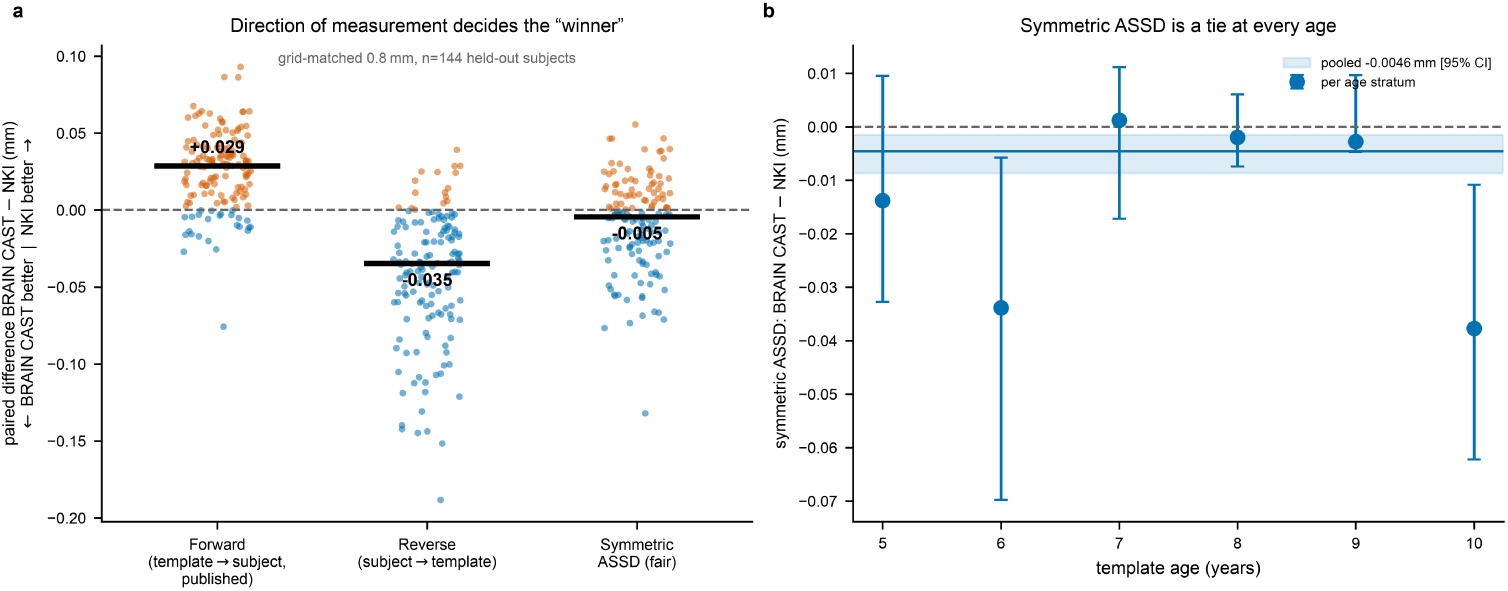
The BRAIN CAST–NKI “winner” is an artifact of measurement direction; the fair symmetric distance is a tie. Grid-matched 0.8 mm, *n* = 144 held-out subjects. **a**, Per-subject paired difference BRAIN CAST−NKI for three formulations of the same interface distance: the one-directional template→subject metric favors NKI (+0.029 mm), the reverse subject→template direction favors BRAIN CAST (−0.035 mm), and the direction-symmetric ASSD is a tie (−0.005 mm; black bars = medians). **b**, The symmetric-ASSD difference with bootstrap 95% CIs is within ±1.5% of a voxel of zero at every age.

**Table 4:** Head-to-head gray–white interface error. Grid-matched (0.8 mm) interface error on held-out HBN subjects (BRAIN CAST/NKI *n* = 144, ages 5–10; Fonov *n* = 209); lower is better. Tabulated medians are the one-directional template-boundary-to-subject-surface distance. On the direction-symmetric ASSD, BRAIN CAST and NKI are equivalent (BRAIN CAST−NKI = −0.005 mm mean / +0.011 mm median, <1.5% of a voxel; Fig. 9), whereas BRAIN CAST decisively beats the broad-age Fonov/NIHPD reference in both measurement directions (209/209 subjects).

| Reference | Full median<br>(mm) | Cortical median<br>(mm) | Paired $\Delta$ vs BRAIN CAST<br>(cortical, mm) |
| --- | --- | --- | --- |
| BRAIN CAST | 0.836 | 0.705 | — |
| NKI | 0.783 | 0.677 | +0.024 (BRAIN CAST higher) |
| Fonov/NIHPD | 1.078 | 0.908 | −0.150 (BRAIN CAST lower) |

Tabulated values are one-directional (template boundary → subject surface) medians; paired BRAIN CAST–NKI = $+0.049$ mm full / $+0.024$ mm cortical ( $\approx 3\text{--}6\%$ of a 0.8 mm voxel). The opposite direction (subject surface → template boundary) favors BRAIN CAST by 0.035 mm cortical (128/144 subjects), and the direction-symmetric ASSD is a statistical tie ( $-0.005$ mm by the mean, $+0.011$ mm by the median). BRAIN CAST is better than Fonov/NIHPD in all 209/209 subjects.

Two points connect this to the validity-vs-reliability thesis below. First, the metric is *SyN-saturated*: in a held-out pilot (*n* = 33 registrations spanning a range of subject–template age mismatches), the interface error is flat in the age mismatch (slope +0.0009 mm yr^−1^, *p* = 0.90), because high-degree-of-freedom nonlinear registration warps away age- and sex-specific shape before the distance is measured. The metric thus cannot reward BRAIN CAST’s age × sex design, and treats a single sharp sex-neutral mean (NKI) as an equally good registration target. Second, that the apparent BRAIN CAST–NKI ranking depends on the measurement *direction*—while the underlying symmetric distance is a tie—is itself an example of reliability-style metric sensitivity (Sec. 3.6): which template appears “best” depends on the summary chosen, not on a stable difference in representativeness. The honest reading is equivalence with NKI on this saturated axis and decisive superiority over both the nearest age-specific competitor (Śanchez/Richards) and the broad-age reference, with BRAIN CAST’s distinctive value—age and sex specificity and per-year granularity—residing on axes this metric is blind to.

### 3.5 Deformation cost and regularity

The age-nine Δ analysis (sex-matched vs. mismatched deformation cost; Supplementary Fig. S1) collapses the full registration matrix onto a single template age. To characterise the age trend directly—and to test it formally—we registered the CSF-normalized intensity image of every held-out subject to BRAIN CAST (Children’s Age- and Sex-specific Templates) spanning a range of ages and measured the clean deformation cost (brain-masked mean displacement of the SyN warp alone, which excludes the subject-specific global transform carried by the affine; see Supplementary Material and Supplementary Fig. S2).

Across 3,190 subject-to-template registrations, cost rises weakly with age mismatch: the median is 2.404 mm at |Δage| < 0.5 y and 2.432 mm at |Δage| ≥ 2 y (Fig. 10a). We deliberately state that difference as a magnitude rather than a verdict, because at this sample size the two part company: +0.028 mm reaches *p* = 0.006 (Mann–Whitney), and 0.028 mm is 3.5% of a 0.8 mm acquisition voxel and 0.06 of the between-subject standard deviation (0.427 mm). The same holds for the trend across the whole matrix—cost increases by 0.025 mm per year of mismatch (*r* = 0.08, *p* = 3 × 10^−6^; Fig. 10b), so that four years of age mismatch, the span of the library’s school-age range, moves the cost by 0.10 mm, a quarter of one standard deviation. Across the entire ±6-year range the binned medians span 0.135 mm, a sixth of a voxel, and the minimum falls at −3 y rather than at the matched age; neither sex shows a matched-age minimum either (Fig. 10c). Age matching is therefore detectable in the deformation-cost domain and immaterial within it, and we report it that way rather than as an effect.

**Fig. 10:**
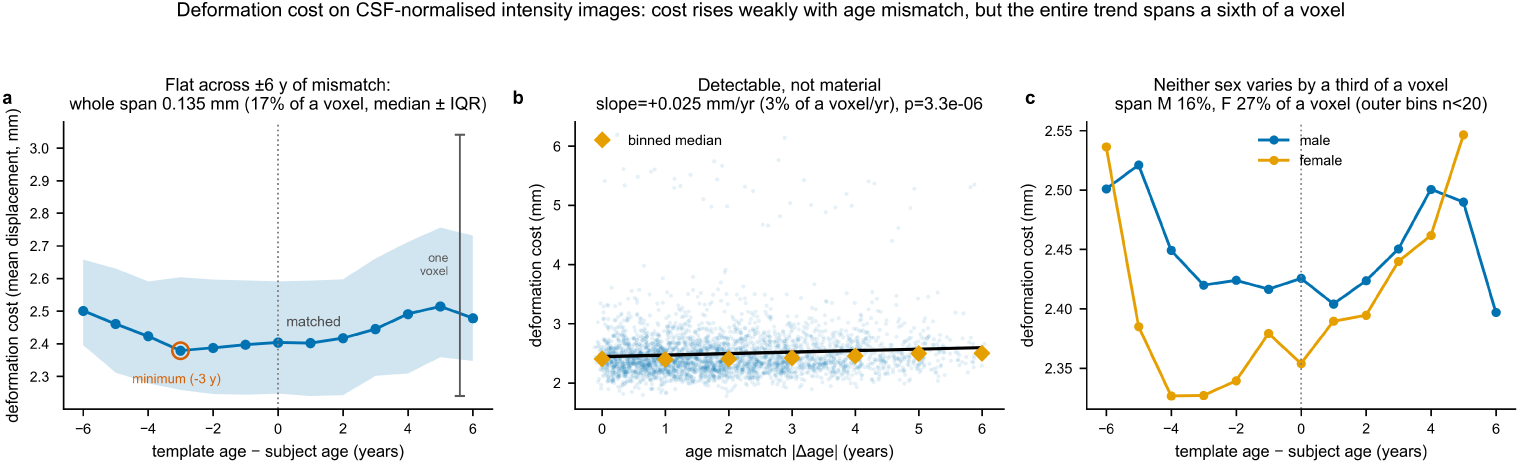
Deformation-cost age trend on CSF-normalized intensity images. Brain-masked mean displacement of the SyN warp across 3,190 subject-to-template registrations. **a**, Cost versus signed age mismatch (template age minus subject age), median ± IQR, with a one-voxel scale bar for reference: the binned medians span 0.135 mm, a sixth of a voxel, across the full ±6-year range, and the minimum falls at −3 y rather than at the matched age. **b**, Cost versus |Δage| with linear fit: +0.025 mm yr^−1^ (3% of a voxel per year, *p* = 3 × 10^−6^)—detectable at *n* = 3,190, immaterial in magnitude. **c**, By sex: neither sex has a matched-age minimum, and the male and female curves span 16% and 27% of a voxel respectively (outermost bins *n <* 20).

Fitting the full linear mixed-effects model of Eq. 1 to this clean displacement metric makes the dominant driver explicit. In the model (*D_i,j_* = *β*_0_ + *β*_1_ Age*_i_* + *β*_2_ Age*_j_* + *β*_3_ |Age*_i_ −* Age*_j_ |* + *β*_4_ SexMatch*_i,j_* + *β*_5_ SubjectSex*_i_* + *u_i_* + *ε_i,j_*, with a per-subject random intercept; *N* = 3,190 registrations across 319 subjects), deformation cost was not associated with subject age (*β*_1_ = −0.002 mm yr^−1^, 95% CI [−0.026, 0.021], *p* = 0.84). Three terms are statistically distinguishable from zero and all three are smaller than a twentieth of a voxel: template age (*β*_2_ = 0.021 mm yr^−1^, 95% CI [0.018, 0.023], *p <* 10^−50^), the absolute subject–template age gap (*β*_3_ = 0.015 mm yr^−1^, 95% CI [0.012, 0.018], *p <* 10^−18^), and sex matching between subject and template (*β*_4_ = −0.014 mm, 95% CI [−0.021, −0.006], *p* = 3 × 10^−4^, i.e. sex-matched registrations are cheaper by 1.7% of a voxel). Subject sex again showed nothing (*β*_5_ = −0.015 mm for male subjects, 95% CI [−0.110, 0.080], *p* = 0.76). Between-subject variance dominates (subject SD = 0.413 mm, residual SD = 0.108 mm; intraclass correlation 0.94): who the child is matters roughly four times more than which template they are registered to. Two of these terms were non-significant in the mask-era fit and *β*_3_ has changed sign, but the practical reading is unchanged and is now better supported— every fixed effect in the model is an order of magnitude below the voxel size, so age and sex matching leave no *materially* meaningful deformation-cost signature once the flexible diffeomorphic warp has acted. A robustness fit pooling the sex-neutral NKI reference templates (coded as sex non-matches) reverses the sign of the SexMatch term (+0.018 mm, *p* = 3 × 10^−6^); that reversal is itself the argument against pooling, since a genuine sex-matching effect cannot depend on whether sex-neutral templates are folded in, and it is therefore not reported as a primary result. The demonstrable sex benefit lives in structural fidelity (Sec. 3.3), not in deformation cost.

This gentle magnitude is expected and is not evidence against age-specificity. School-age brains are already ∼90% of adult size, and high-degree-of-freedom diffeomorphic registration absorbs most residual age and shape differences, so a flexible warp can fit a mildly mismatched template at almost the same cost. Deformation cost is therefore a *reliability* measure that is confounded by deformation smoothness— an over-regularised or over-smooth template can register at low cost while being anatomically wrong (the “reliable but wrong” trap). The value of an age- and sex-matched template is better captured by downstream structural *bias*, which we quantify directly with the gray–white interface metric (next section); there, matching produces a sub-voxel improvement that the deformation-cost metric, by construction, cannot reveal.

#### Tissue-specific deformation regularity

To localize the deformation within tissue compartments, we extended the age-nine Jacobian-regularity analysis (Supplementary Fig. S3) to gray matter, white matter, and CSF using template-space FAST segmentations, over the matched-template SyN warps of 81 held-out subjects spanning ages 5–12 (Fig. 11). As in Sec. 2.6, the Jacobian is taken from the SyN warp field alone, with the affine excluded, so these values are directly comparable with the construction-warp Jacobians and not with composed ones. The mean log-Jacobian was near zero in every tissue (gray matter −0.03, white matter −0.03, CSF +0.05), indicating no systematic volumetric bias: cortical and white matter are marginally compressed and CSF marginally expanded when a subject is normalized to the template, consistent with the small inward gray–white bias reported above. Local deformation magnitude (the SD of the log-Jacobian) was largest in gray matter (0.25) and smaller in white matter (0.22) and CSF (0.20), as expected given the greater shape variability of the cortical ribbon. Critically, the fraction of non-diffeomorphic voxels (*J ≤* 0) was 0% in all three tissues, confirming that the registrations preserve topology everywhere and fold or tear no tissue compartment.

**Fig. 11:**
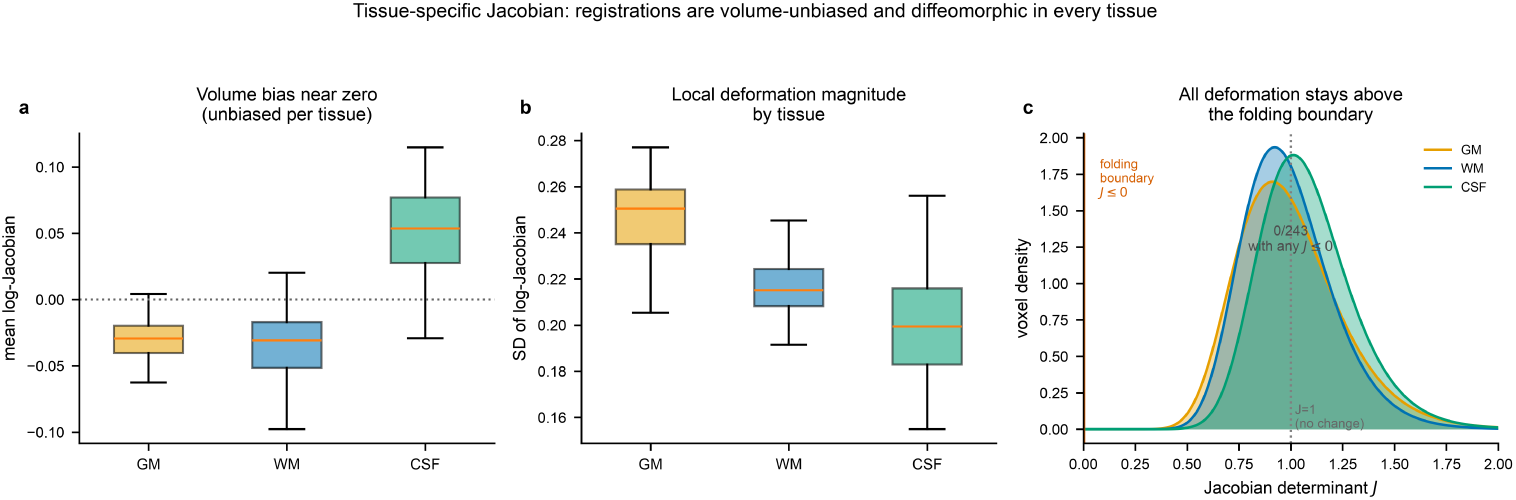
Tissue-specific Jacobian. (81 held-out subjects, matched-template SyN). **a**, Mean log-Jacobian per tissue is near zero (volume-unbiased). **b**, SD of the log-Jacobian is largest in gray matter (most local deformation). **c**, Voxelwise Jacobian-determinant distribution per tissue (modelled from each subject’s log-Jacobian mean and SD, voxel-count weighted): the mass clusters at *J ≈* 1 and stays well clear of the folding boundary (*J ≤* 0). No voxel was non-diffeomorphic in any subject or tissue (0/243 subject–tissue measurements, computed directly from the determinant maps).

### 3.6 Validity versus reliability and pipeline ablation

The preceding results motivate an explicit distinction between two families of template-quality metric (Table 5). *Reliability* metrics are image-based—edge sharpness, gray–white contrast, fractal dimension, and the regularity/cost of the deformation field. They are necessary but not sufficient: a template can score well on them while poorly representing anatomy (the “reliable but wrong” failure mode). On these metrics BRAIN CAST is comparable to the NKI reference (edge sharpness is not comparable across intensity-normalization conventions; Weber contrast is similar; deformation cost is comparable, 2.42 vs 2.38 mm—a paired comparison of the same 319 subjects registered to both families under identical inputs and settings, differing by 0.044 mm, 5.5% of a voxel), with the one scale-invariant advantage being higher white-matter fractal dimension (2.34 vs 2.30; a modest but unanimous +0.041 across the eleven shared ages that widens under a matched-resolution control, +0.058, 11/11). *Validity* —the downstream structural bias a template induces—is the property that ultimately matters, and it is where the age- and sex-specific construction pays off: the gray–white interface error is uniformly sub-voxel (1.05–1.26 mm across strata) and is not improved by higher image detail: across the sixteen strata its correlation with white-matter fractal dimension is weakly *positive* (Fig. 12; Pearson *r* = +0.44, *p* = 0.09; Spearman *ρ* = +0.55, *p* = 0.03; *n* = 16), so the more detailed templates are, if anything, marginally less accurate. In other words, image-quality metrics do not, by themselves, establish representativeness—and what association they do carry runs the wrong way to serve as a quality proxy; the structural-bias measurement does. We therefore foreground validity over sharpness throughout. The head-to-head benchmark above reinforces this point from a different angle: on the SyN-saturated interface metric BRAIN CAST and NKI are equivalent, and the apparent ranking between them depends on the *direction* in which the distance is measured—the one-sided boundary-to-surface form favors NKI while the direction-symmetric ASSD is a tie (Table 4, Fig. 9)—a metric-sensitivity effect of exactly the reliability kind, which is why we do not rest BRAIN CAST’s contribution on this single axis.

**Fig. 12:**
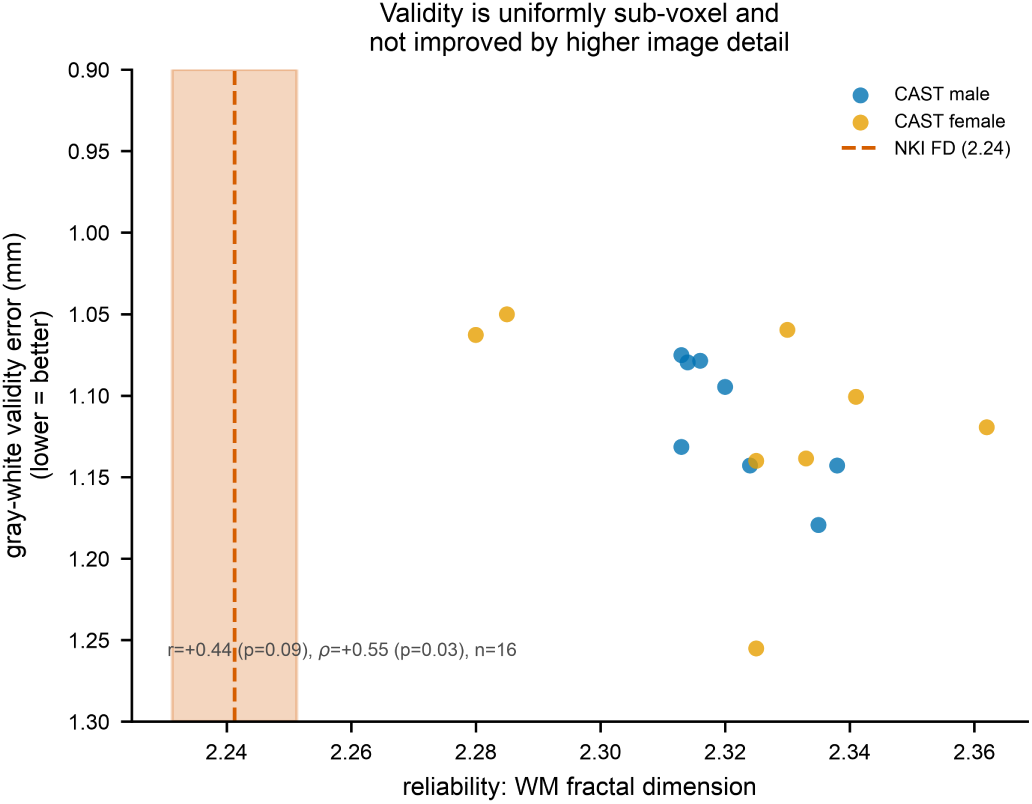
Validity versus reliability. Per-stratum gray–white validity error against white-matter fractal dimension (a reliability/detail metric), all 16 strata: validity is uniformly sub-voxel and is not improved by higher image detail—the association is weakly positive, i.e. in the unhelpful direction, and does not reach significance by Pearson (*r* = +0.44, *p* = 0.09; Spearman *ρ* = +0.55, *p* = 0.03). The NKI reference fractal dimension is marked.

**Table 5:**
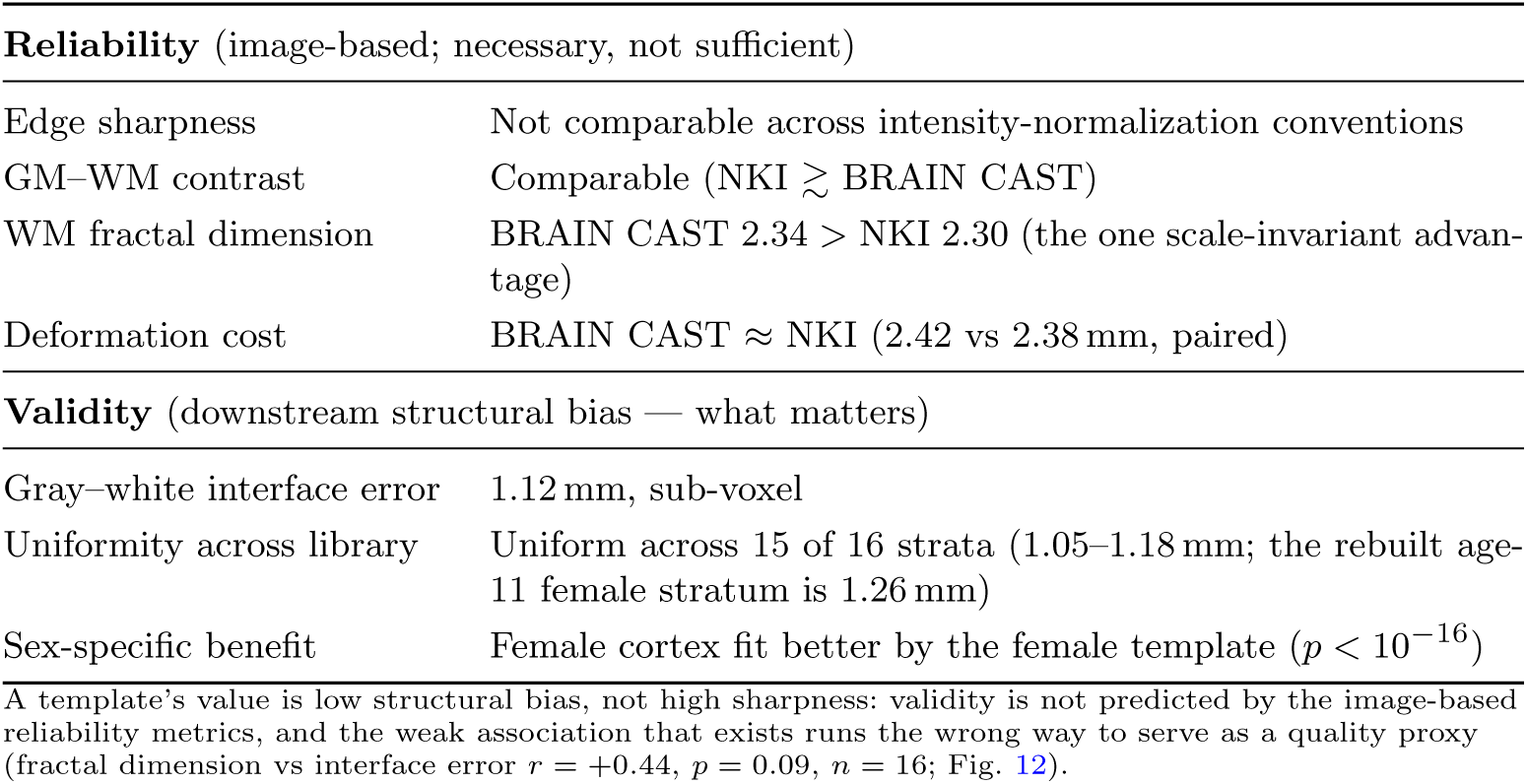
Reliability versus validity metrics. Image-based reliability metrics are necessary but not sufficient and do not, by themselves, separate templates; the downstream structural-bias (validity) metric is the property that establishes representativeness.

#### Pipeline ablation: visual characteristics across pipeline variants

We evaluated how successive pipeline modifications influenced template appearance by comparing templates generated with the original pipeline, an intermediate pipeline incorporating preprocessing improvements, and the final pipeline integrating both pre-processing and registration refinements (Fig. 13). We iteratively tuned these steps using MRIQC-derived sharpness and smoothness indicators (Tenengrad, entropy-focus criterion (EFC) and whole-brain FWHM) alongside tissue-contrast ratios (WM/GM, GM/CSF and WM/CSF; Supplementary Table S1). In the qualitative comparison, preprocessing updates improved tissue contrast and boundary clarity relative to the original workflow, and the final pipeline further improved edge definition and overall visual fidelity. Quantitatively, the final pipeline achieves the lowest global smoothness of any variant (whole-brain FWHM 3.82 vox, against 4.00 and 5.06 for the original and intermediate pipelines), but the remaining image metrics do *not* consistently favour it: its entropy focus criterion is higher than either earlier variant (0.341 vs 0.234 and 0.256). We report these values in Supplementary Table S1 rather than the main text because they cannot bear the interpretive weight of a quality comparison. Successive pipeline revisions deliberately changed the intensity-normalization convention—mean white-matter intensity is 17.1, 15.6 and 8.3 in the original, intermediate and final templates, and 7110 in the raw-scanner-unit NKI reference—so any gradient-energy measure, which scales as the square of intensity, is not comparable across these columns at all, and we therefore omit it. This is exactly the reliability/validity dissociation documented above: image-based sharpness and contrast metrics do not identify the better template. Our representativeness evidence rests entirely on the downstream structural-fidelity benchmark, which is invariant to intensity convention.

**Fig. 13:**
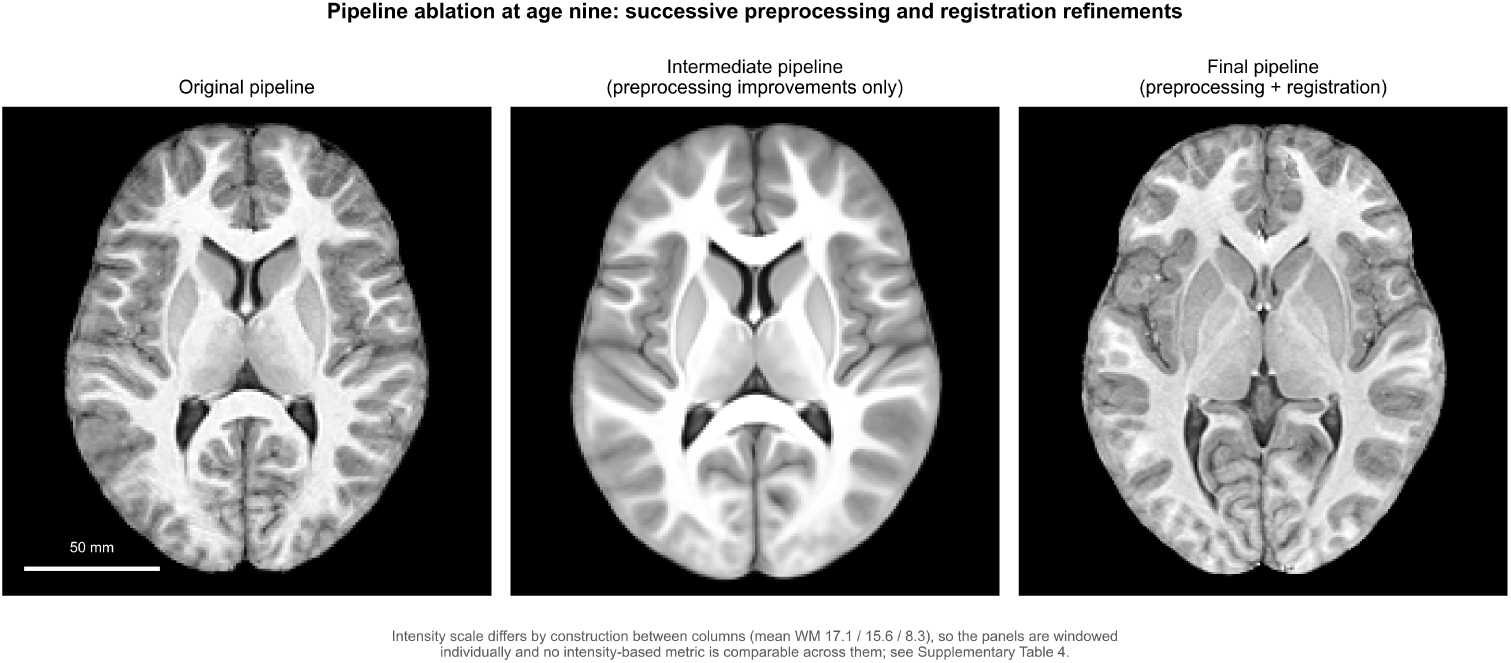
Effects of pipeline improvements on template visual characteristics. Matched axial sections through three age-nine templates built with the original pipeline, an intermediate pipeline incorporating the preprocessing improvements only, and the final pipeline with both preprocessing and registration improvements. The final pipeline yields clearer edge definition and visibly more cortical detail; the intermediate template is the smoothest of the three, consistent with its whole-brain FWHM of 5.06 vox against 4.00 and 3.82 (Supplementary Table S1). Panels are reoriented to a common anatomical frame, centred on each template’s own brain centroid and drawn on a shared 180 mm field of view at true physical scale, so apparent size differences are anatomical rather than an artifact of the differing voxel sizes (1.0, 1.0 and 0.8 mm). Intensity windowing is per-panel because the three pipelines use different intensity-normalization conventions (mean WM 17.1, 15.6 and 8.3), which is also why no intensity-based metric is comparable across them.

## 4 Discussion

We have developed and demonstrated a reproducible, open-source pipeline that leverages quantitative quality control to generate a high-fidelity library of age- and sex-specific pediatric brain templates for ages 5 through 18. By processing over 2,400 aggregated T1-weighted images from the Healthy Brain Network cohort [1] (1,272 contributing after quality screening), we constructed templates in fine-grained, one-year age intervals, stratified by sex. The methodological framework integrates MRIQC for iterative preprocessing refinement [3], DeepBET for robust skull stripping [4], and the well-established ANTs symmetric diffeomorphic normalization (SyN) algorithm for groupwise registration [5, 24]. The resulting templates and the registration fields used to create them were evaluated for anatomical sharpness, tissue contrast, deformation cost, mathematical regularity, and—most directly—the downstream gray–white structural fidelity they induce in held-out subjects, demonstrating measurable improvements over less refined approaches and providing a quantitative basis for their use in future neuroimaging studies.

Our evaluation strategy distinguishes template *reliability* from template *validity*. Image-based metrics—edge sharpness, tissue contrast, and the cost and regularity of the deformation field—are reliability measures: they are necessary but not sufficient, because a template can register at low cost while poorly representing anatomy (the “reliable but wrong” failure mode), and deformation cost in particular is confounded by deformation smoothness. We therefore treated the downstream structural bias—the gray–white interface error remaining after a held-out subject is normalized to the matched template—as the primary index of template quality, and found it to be uniformly sub-voxel (≈1.1 mm across strata), the operative definition of representativeness for morphometric and surface-based analyses. The analysis of Jacobian determinants provides a complementary measure of the transformation’s regularity and physical plausibility: by confirming that all Jacobian determinants were positive, we ensure the transformations are diffeomorphic and preserve topology, meaning they do not create artificial folds or tears in the brain anatomy [5]. Deformation cost, by contrast, proved an insensitive index of template fit—a flexible diffeomorphic warp absorbs most age and shape mismatch—which is why we foreground structural fidelity rather than registration cost throughout.

The construction of separate templates for males and females allowed us to quantify the utility of sex-specific references during pediatric development, and our results locate that utility in structure rather than in registration cost. At the gray–white interface, the genuine evidence is a sex × subject-sex *interaction* (mixed-effects *p* = 7 × 10^−3^) rather than a uniform matched-sex advantage: held-out female subjects fit the female template significantly better than the male template (by 0.049 mm cortical, 0.129 mm full; Wilcoxon *p* = 1*imes*10^−17^, better in 95/99 subjects), whereas male subjects show a reversal, fitting the female template ≈ 0.038 mm better (*p* = 2*imes*10^−28^). We attribute this reversal to the female template being an easier registration target for all subjects (by ≈ 0.042 mm)—a confound we disclose explicitly—so that the substantive signal is the interaction, in which female cortical shape is fit better by its own template, rather than a blanket matched-sex rule. A single sex-neutral reference cannot provide sex specificity at all. Deformation cost, by contrast, resolves a sex-matching term of only 0.014 mm—detectable at *n* = 3,190 but under a fiftieth of a voxel, and under a fifth of the structural effect—because a high–degree-of-freedom diffeomorphic warp largely equalizes it. The sex-specific benefit is therefore a structural-fit difference measurable on cortical anatomy, consistent with known periods of sexually dimorphic neurodevelopment [12–14, 53]; it is a measure of template fit, not a direct biological statement on the magnitude or nature of sex differences in brain structure. These templates provide a tool to reduce sex-related structural bias, thereby enabling more sensitive downstream analyses.

How accurate a template must be depends on the application, and our measured fidelity can be read against those requirements. The most demanding uses—voxel- and surface-based morphometry, cortical-thickness estimation—require boundary accuracy at the millimetre scale, because blurred or displaced gray–white boundaries directly bias thickness and volume measurements and can collapse shallow sulci; the matched template’s sub-voxel (≈1.1 mm, 84–89% within 2 mm) interface error sits comfortably within this tolerance. Less geometrically demanding applications have far larger budgets: in EEG/MEG source localization the forward head model tolerates errors on the order of a centimetre, and adult or age-mismatched head models are reported to introduce 10–20 mm localization errors [17, 18], so an age- and sex-matched pediatric template with millimetre-scale structural fidelity removes template mismatch as a limiting factor for that use case. Characterizing how the residual error propagates is therefore important: the gray–white error we observe is small, spatially diffuse, and slightly biased inward (the template boundary lies ∼0.1–0.2 mm interior to the subject’s), so for absolute morphometry it imposes a minor systematic offset, whereas for group analyses that register all subjects to a common template it largely cancels. The dominant avoidable error—sulcal collapse and boundary blurring incurred by normalizing to a mismatched reference—is precisely what age- and sex-matching mitigates, keeping the interface error sub-voxel. In practice these templates are therefore most valuable for structural and surface analyses of school-age children aged 5–18 when matched on age and sex (with female cortex showing the clearest cortical-interface benefit from its own template, per the sex × subject-sex interaction above), while the requirements are easily met for coarser geometric applications; their main current limits are cross-population and cross-scanner generalization and the smaller late-adolescent strata.

This study has several important limitations. Our evaluation combined image-quality and deformation-field metrics with a more direct measure of downstream gray–white structural fidelity; nonetheless, none of these constitutes validation against ground-truth anatomical segmentations, which are unavailable for living subjects. In particular, the reported ≈1.12 mm gray–white interface error should be read as an upper bound rather than a pure template–subject discrepancy: the reference surface is itself a FreeSurfer reconstruction, whose white-surface placement error against histology and manual tracings is on the order of 0.3–0.5 mm, so a comparable component of the measured 1.12 mm reflects the surface-extraction floor rather than template inaccuracy alone. The structural-validity analysis was conducted on held-out subjects aged 5–12—the developmental window of primary interest for this library—so the cross-sex interaction and its trajectory into later adolescence remain to be characterized across the full age range; extension to adolescents (ages 13–18, for whom both FreeSurfer reconstructions and centered images are available in the test set) is feasible but the upper-age strata are thinly and unevenly populated (down to *n* = 2 at age 18) and would not be evenly sampled across the adolescent range. More generally, several late-adolescent and minority-sex strata in Table 2 are thinly populated (e.g. 13 female subjects at age 15, 11 at age 18), and templates built from such small cohorts are correspondingly noisier; these strata should be used with caution. The sex-specific benefit should likewise be calibrated rather than over-read: it manifests as a sex × subject-sex interaction in which female cortex is fit ≈ 0.049 mm better by the female template, while males show a reversal driven partly by the female template being an easier target for everyone (≈ 0.042 mm); it is therefore not a uniform matched-sex advantage, and it must be weighed against the cost of sex stratification, which roughly halves the number of subjects contributing to each template and thereby increases per-template sampling variability; sex-specific construction is therefore most justified where the female cortical-fit benefit and the ability to represent sex-dimorphic anatomy (which a sex-neutral reference cannot) outweigh the loss of statistical stability from the smaller per-template *n*. The templates are derived from a large but specific cohort (HBN) and their generalizability to other populations, scanners, or clinical cohorts remains to be established; BRAIN CAST is built from children characterized as typically developing within a transdiagnostic recruitment frame (HBN is a community sample enriched for mental-health and learning concerns), so “typically developing” here denotes the absence of gross structural abnormality on screening rather than a strictly healthy or clinically vetted population, and applicability to clinical or atypical anatomy should not be assumed. Demographic and developmental moderators that we did not model— pubertal stage (which dissociates from chronological age, especially in the 9–14-year range) and socioeconomic status—may influence both anatomy and the appropriateness of one-year chronological-age binning; finer or puberty-aware binning could refine age specificity but would further reduce per-bin sample sizes. We also note that the present templates and validity analysis are whole-brain and do not include extracerebral tissue (skull, scalp); applications that depend on a complete head model—notably EEG/MEG forward modelling and source localization, where geometric accuracy of the skull and scalp compartments governs localization error [17, 18]—would require extending the construction to whole-head, multi-compartment templates, which is beyond the present scope. Furthermore, our CSF-anchored intensity normalization, while principled, means that the resulting template intensities are dependent on this specific scheme. The templates are also cross-sectional and do not capture longitudinal changes within individuals.

Future work can extend this methodology in several key directions. The linear mixed-effects model reported here quantifies the deformation-cost trends; extending it with nonlinear age terms and additional regularity covariates could further characterize the observed age- and sex-specific effects. The inclusion of additional regularity metrics, such as measures of shear and strain, could provide a more complete picture of the deformation fields. The framework presented here is extensible to broader template families, including the incorporation of multimodal data (e.g., T2-weighted, DTI) to create richer, tissue-specific atlases, or the construction of longitudinal templates from serial scan data. Amortized, learning-based conditional-atlas approaches that generate continuous age- and sex-conditioned templates [33] are a complementary future direction to the discrete one-year, sex-stratified construction used here. Finally, external validation of these templates on independent pediatric datasets is a crucial next step to confirm their robustness and broad utility.

In conclusion, this work provides a robust, extensible, and openly available resource for the pediatric neuroimaging community. By combining rigorous quality control with advanced registration techniques, we have produced a set of age- and sex-specific templates that offer improved anatomical fidelity and reduce normalization bias. These tools provide a more accurate reference for studying brain development and will support more precise and sensitive analyses of structural MRI data in children and adolescents.

## Acknowledgements

Computational resources and services were provided by the Research Computing Data Core at the University of Houston (https://www.uh.edu/research/rcdc/index.php). This work was supported in part by the NSF IUCRC Building Reliable Advances and Innovation in Neurotechnology (BRAIN) Center (https://brain-d10.egr.uh.edu, Award #2137255).

## Data availability

The BRAIN CAST template library is derived from the Healthy Brain Network (HBN) and is not a Child Mind Institute product; the underlying subject-level T1-weighted images (ages 5–18) are distributed by HBN through the International Neuroimaging Data-Sharing Initiative [1]. The constructed BRAIN CAST templates are openly available from Zenodo under the concept DOI 10.5281/zenodo.21491758, which always resolves to the most recent version of the deposit, under a CC-BY 4.0 license. The full file layout, template naming conventions, and per-stratum cohort composition are documented in the companion data descriptor (Hu & Contreras-Vidal, in review, *Scientific Data*). The dataset accession is 10.5281/zenodo.21491758.

## Code availability

The full pipeline code, including preprocessing scripts, container definitions, and analysis notebooks, is openly available on GitHub at https://github.com/bifeitang/cast-pipeline and archived on Zenodo under the concept DOI 10.5281/zenodo.21482327, which always resolves to the most recent release, under the MIT license. The as-run Docker image (bifeitangmac/mri_template_env_matlab) is hosted on DockerHub for direct use (https://hub.docker.com/r/bifeitangmac/mritemplate_env_matlab), and an Apptainer/Singularity recipe is provided for HPC environments. The DeepBET tool used is open-source (https://github.com/wwu-mmll/DeepBET). Custom modifications or parameter configurations for ANTs are documented in the repository. Users can reproduce the template construction and all reported analyses using the code and instructions in the README. Parameters and random seeds are fixed in the code for exact reproducibility.

## Ethics declaration

All data were obtained from the Healthy Brain Network (HBN), which was approved by the Chesapeake Institutional Review Board (now Advarra IRB) and conducted in accordance with the Declaration of Helsinki. Written informed consent was obtained from participants aged 18 or older, and written informed consent from a legal guardian together with written assent from the child was obtained for participants under 18 [1]. The present work is a secondary analysis of these de-identified, openly distributed data and introduces no new human-subjects data collection. Use of the de-identified, openly released HBN data for the present study did not require additional institutional review board approval at the University of Houston. The released templates are group-level averages containing no subject-identifiable data.

## Author contributions

Y.H. designed and implemented the preprocessing and template construction pipeline, performed all computational experiments and quality control, and drafted the manuscript. J.L.C-V. supervised the project, contributed to study design and interpretation, and revised the manuscript. Both authors read and approved the final manuscript.

## Supplementary Material

### Pipeline ablation: image metrics across pipeline variants

Table S1 reports the image-quality metrics recomputed for the four template variants compared in Sec. 3.6: the NKI age-nine reference, and the templates produced by the original, intermediate and final BRAIN CAST pipelines. All values were regenerated with a single documented script (pipeline_ablation_metrics.py, archived with the pipeline release) applied to the four template volumes; tissue classes come from FSL FAST (-t 1 -n 3) on the brain-masked volume, sampled at pve *>* 0.9 so that partial-volume voxels do not contaminate the class means—the same segmentation procedure used for the template morphometry in Sec. 3.5. EFC follows Atkinson’s entropy focus criterion as implemented in MRIQC; whole-brain FWHM is the mean of the three axis estimates from the Forman finite-difference estimator, in voxels.

We deliberately omit gradient-energy (Tenengrad) measures. The mean white-matter intensity row makes the reason explicit: successive pipeline revisions changed the intensity-normalization convention by roughly a factor of two, and the NKI reference is in raw scanner units some three orders of magnitude larger again. Any gradient-energy statistic scales as the square of intensity, so such a comparison across these four columns measures the normalization convention rather than image sharpness. For the same reason the tissue-contrast ratios, though scale-invariant under multiplicative rescaling and therefore reported here, should be read as descriptive rather than as a ranking. The table shows that these image metrics do not consistently identify the final pipeline as best—only whole-brain FWHM does—which is the empirical basis for the validity-over-reliability position taken throughout the main text.

**Table S1:** Image-quality metrics across pipeline variants (recomputed). Lower EFC and lower whole-brain FWHM indicate less blur and lower global smoothness. Tissue-contrast ratios are ratios of mean intensity between FAST tissue classes. The final row is not a quality metric: it is given so the reader can see that the four volumes occupy different intensity scales, which is why no gradient-energy metric is reported.

| Metric | NKI age 9<br>reference | Original<br>pipeline | Intermediate<br>pipeline | Final<br>(released) |
| --- | --- | --- | --- | --- |
| EFC ↓ | 0.378 | 0.234 | 0.256 | 0.341 |
| FWHM (vox) ↓ | 4.31 | 4.00 | 5.06 | **3.82** |
| WM/GM | 1.551 | 1.574 | 1.588 | 1.429 |
| GM/CSF | 3.618 | 4.519 | 5.250 | 3.774 |
| WM/CSF | 5.611 | 7.111 | 8.335 | 5.394 |
| mean WM intensity (*scale only*) | 7109.7 | 17.13 | 15.62 | 8.32 |

### Sex-stratified deformation-cost differences

For each subject, we quantified the difference in normalized deformation cost Δ when registering to the sex-matched versus mismatched age-nine template (Fig. S1). Positive Δ values indicate a lower deformation cost for the sex-matched template. Waterfall plots show that a majority of *male* subjects have positive Δ (69%, *n* = 204), whereas female subjects split almost evenly (45%, *n* = 115)—the same sex asymmetry the structural-fidelity analysis finds, and not a uniform matched-sex advantage. When stratified by age, Δ does *not* peak at the template age: for males it is largest at age 10 (+0.0009) rather than age 9 (+0.0006) and is essentially flat across ages 5–12, and for females it is not consistently signed at any age, reaching −0.0004 at age 11. Any age structure here is within the noise of thinly populated per-age bins (*n* = 5–41). This matched-template advantage is modest and, as the translation-artifact-free analysis in the main text shows, is not statistically robust in the deformation-cost domain; the demonstrable sex benefit instead lives in structural fidelity, as shown by the gray– white interface analysis above. In the main text we quantify the underlying age trend with explicit statistics on the full age×age matrix using this translation-artifact-free cost metric (Fig. 10).

**Fig. S1:**
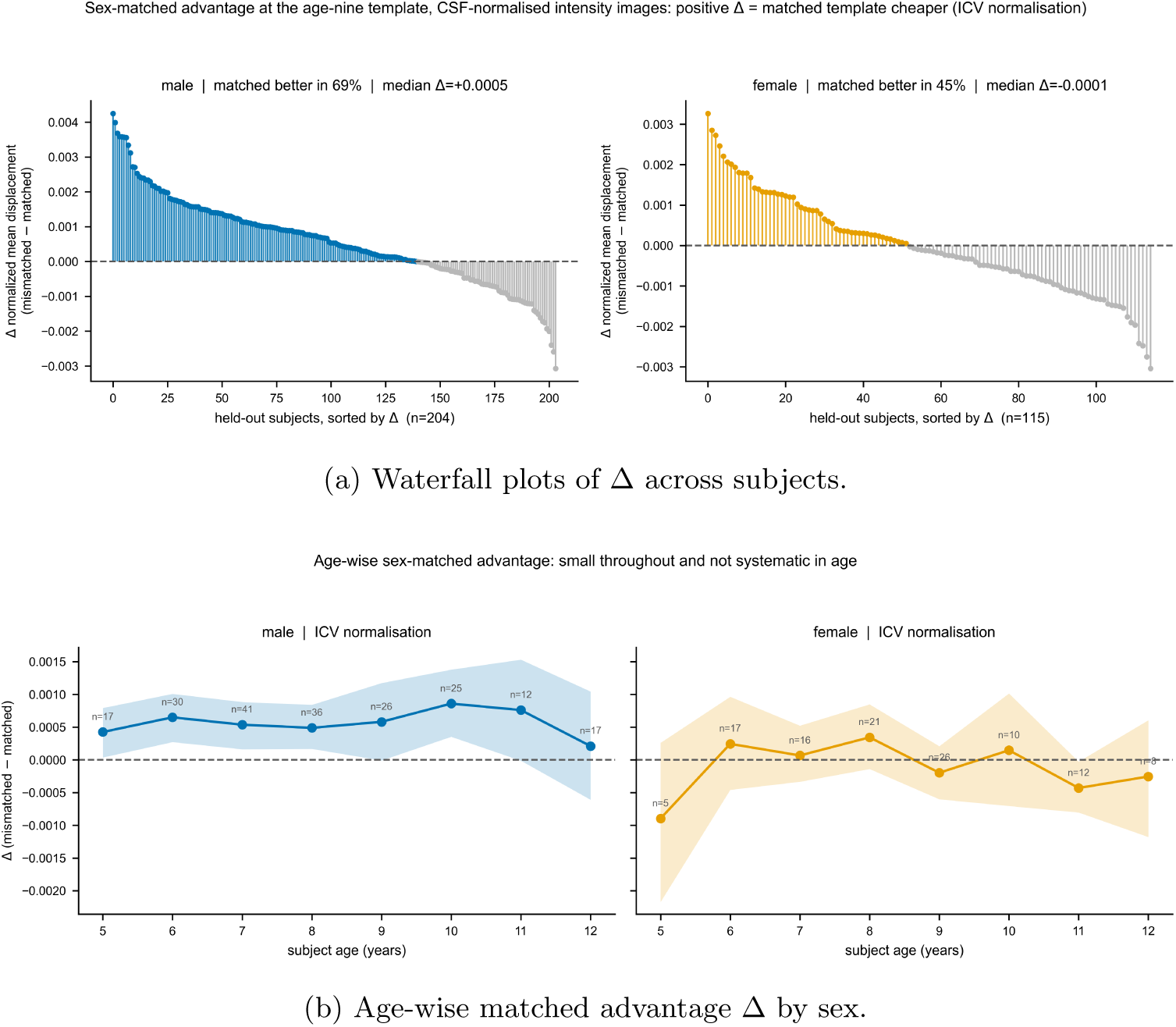
Normalized deformation-cost differences by sex and age. a, Waterfall plots of Δ (mismatched minus sex-matched normalized deformation cost, intracranial-volume normalization) across the pediatric test set; positive values indicate lower cost when using the sex-matched template. The matched template is cheaper for 69% of male subjects (*n* = 204, median Δ = +0.0005) but only 45% of female subjects (*n* = 115, median Δ = −0.0001). b, Age-wise matched advantage Δ by sex; the effect is small throughout and, contrary to the expectation that it would peak at the template age, is not systematic in age. Bands are 95% bootstrap confidence intervals of the mean. Both panels use the intracranial-volume normalization only: the diagonal scheme was not carried by the intensity re-registration, and plotting it from the superseded run would mix two generations of the metric within one figure.

### Why the affine component is excluded

The deformation-cost analyses in the main text measure the SyN warp alone. To quantify what excluding the affine leaves out, we re-registered 647 held-out subject × template pairs spanning all ten templates and measured both components of the same registration (Supplementary Fig. S2). Including the affine multiplies the cost by 2.8 (median 6.81 vs 2.42 mm), and the excess it contributes is not a fixed offset that could simply be subtracted: it has a median of 4.23 mm but ranges from 0.6 to 22.9 mm across registrations (coefficient of variation 0.58), while being statistically flat in subject age (slope +0.07 mm yr^−1^, *p* = 0.21) and near-identical in median across template ages (3.97–4.32 mm). Critically, the composite correlates only weakly with the SyN cost it would replace (*r* = +0.27), so it is largely orthogonal to the anatomical mismatch the metric exists to capture. Retaining the affine would therefore add a large subject-specific nuisance term to every comparison without adding information about template fit, which is why we report the SyN component throughout. The SyN-only median here, 2.416 mm over an independently drawn sample, reproduces the 2.415 mm of the main analysis (Sec. 3.5).

**Fig. S2:**
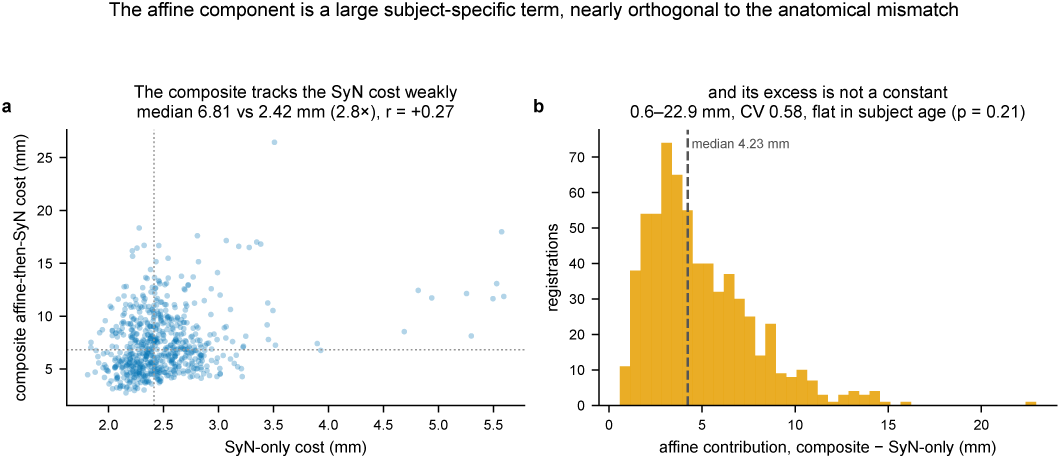
The affine component is a large subject-specific term, nearly orthogonal to the anatomical mismatch. 647 held-out subject × template registrations spanning all ten templates, with both components measured on the *same* registration. **a**, Composite affine◦SyN cost against SyN-only cost: the composite is 2.8× larger (median 6.81 vs 2.42 mm) but tracks it only weakly (*r* = +0.27). **b**, The affine’s contribution, composite minus SyN-only. It is not a constant that could be subtracted: median 4.23 mm over a 0.6–22.9 mm range (coefficient of variation 0.58), flat in subject age (*p* = 0.21) and near-identical in median across template ages. A large nuisance term that carries little information about template fit is the reason the SyN component is reported throughout.

### Jacobian regularity at the age-nine templates

To evaluate how warp regularity behaves under age mismatch, we examined log-Jacobian statistics for the held-out test set registered to the age-nine templates (Supplementary Fig. S3): 319 subjects aged 5–12, each registered to both the male and the female age-nine template, giving 638 registrations. Unlike the construction-warp and tissue Jacobians above, these are computed on the *composed* affine ◦ SyN field, so they are not comparable with the −0.032 of Sec. 2.6 and we do not compare them; the affine’s scale term is precisely what makes the two quantities different. Read within that convention, the mean log *J* was −0.011 pooled over both templates, but the pooled value is the mean of two curves offset from each other by a constant +0.113 at every subject age, and that offset is template size rather than warp behaviour: the age-nine male template is 1558 ml in intracranial volume against the female’s 1389 ml, and the log of that ratio is +0.115. Within each template, mean log *J* rises by ≈ 0.08 across ages 5–12, so younger subjects are compressed onto a nine-year-old target and older ones expanded—the direction anatomy predicts. The within-warp standard deviation of log *J* was nearly flat in subject age, spanning only 0.227–0.251, and reached its minimum at age 10 rather than at the template age. Warp regularity therefore does not degrade as subject age departs from the template age, even though registration cost rises slightly (Sec. 3.5); the two are separate properties of the deformation. No voxel in any of the 638 registrations was non-diffeomorphic. The two normalization schemes yield identical Jacobian statistics, as they must: normalization rescales the displacement metrics but leaves the warp itself unchanged.

**Fig. S3:**
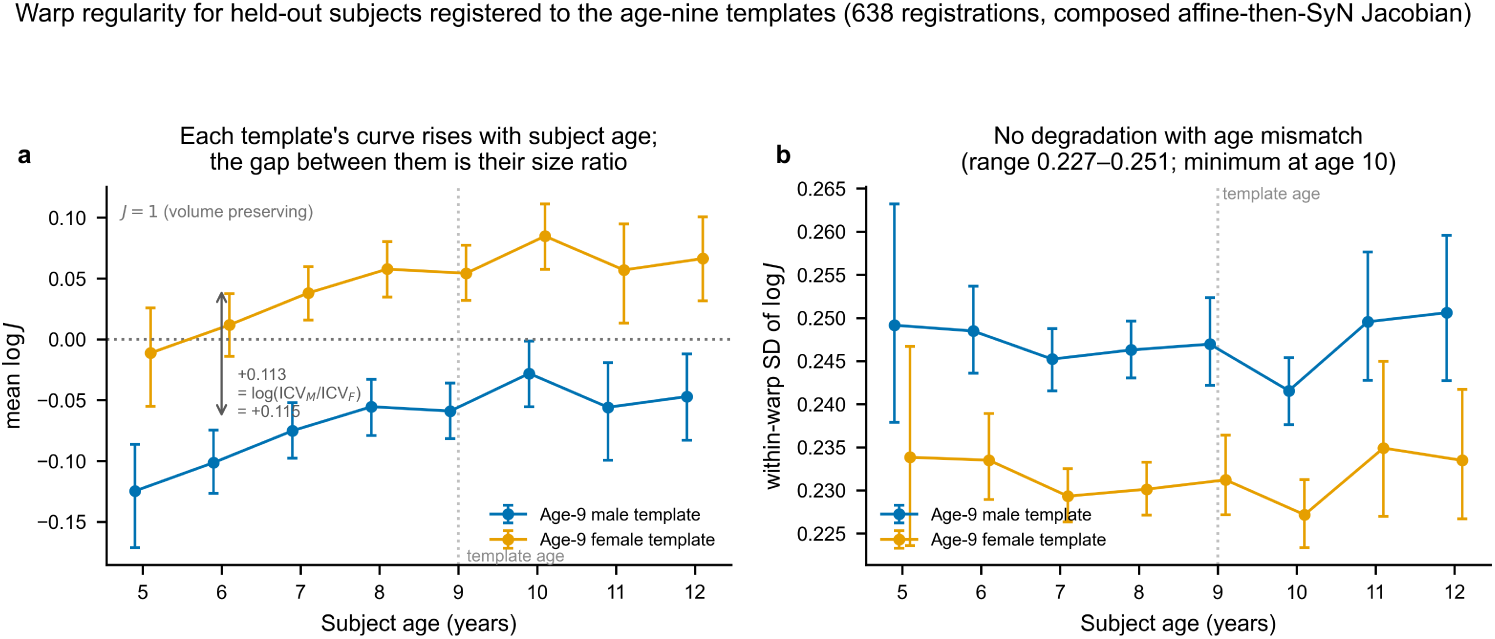
Warp regularity for held-out subjects registered to the age-nine templates. 319 held-out subjects aged 5–12, each registered to both age-nine templates (638 registrations), on CSF-normalized intensity images. Jacobians here come from the *composed* affine ◦ SyN field and are therefore not comparable with the SyN-only construction-warp Jacobians of Fig. 3. **(a)** Mean log *J* by subject age. Each template’s curve rises by ≈ 0.08 across ages 5–12—younger subjects compressed onto a nine-year-old target, older ones expanded—and the constant +0.113 gap between the two curves is the templates’ intracranial-volume ratio (log(1558/1389) = +0.115), not a property of the warps. **(b)** Within-warp standard deviation of log *J*, the magnitude of local volume change, spans only 0.227–0.251 and reaches its minimum at age 10 rather than at the template age (dotted line): regularity does not degrade with age mismatch. Bars are 95% bootstrap confidence intervals. No voxel was non-diffeomorphic in any registration. Intracranial-volume and diagonal normalization give identical Jacobian statistics, so a single series is plotted per template.

